# Global transcriptional response of *Methylorubrum extorquens* to formaldehyde stress expands the role of EfgA and is distinct from antibiotic translational inhibition

**DOI:** 10.1101/2021.01.07.425672

**Authors:** Jannell V. Bazurto, Siavash Riazi, Simon D’Alton, Daniel E. Deatherage, Eric L. Bruger, Jeffrey E. Barrick, Christopher J. Marx

**Affiliations:** Department of Biological Sciences, University of Idaho, Moscow, ID, USA; Institute for Modeling Collaboration and Innovation, University of Idaho, Moscow, ID, USA; Institute for Bioinformatics and Evolutionary Studies, University of Idaho, Moscow, ID, USA; Department of Molecular Biosciences, Center for Systems and Synthetic Biology, The University of Texas at Austin, Austin, Texas, USA; Department of Plant and Microbial Biology, University of Minnesota, Twin Cities, MN, USA; Microbial and Plant Genomics Institute, University of Minnesota, Twin Cities, MN USA; Biotechnology Institute, University of Minnesota, Twin Cities, MN USA

**Keywords:** Formaldehyde, stress response, translation inhibition, kanamycin, methylotrophy, Enhanced formaldehyde growth EfgA, EfgB, proteotoxicity

## Abstract

The potency and indiscriminate nature of formaldehyde reactivity upon biological molecules make it a universal stressor. However, some organisms such as *Methylorubrum extorquens* possess means to rapidly and effectively mitigate formaldehyde-induced damage. EfgA is a recently identified formaldehyde sensor predicted to halt translation in response to elevated formaldehyde as a means to protect cells. Herein, we investigate growth and changes in gene expression to understand how *M. extorquens* responds to formaldehyde with and without the EfgA-formaldehyde-mediated translational response, and how this mechanism compares to antibiotic-mediated translation inhibition. These distinct mechanisms of translation inhibition have notable differences: they each involve different specific players and in addition, formaldehyde also acts as a general, multi-target stressor and a potential carbon source. We present findings demonstrating that in addition to its characterized impact on translation, functional EfgA allows for a rapid and robust transcriptional response to formaldehyde and that removal of EfgA leads to heightened proteotoxic and genotoxic stress in the presence of increased formaldehyde levels. We also found that many downstream consequences of translation inhibition were shared by EfgA-formaldehyde- and kanamycin-mediated translation inhibition. Our work uncovered additional layers of regulatory control enacted by functional EfgA upon experiencing formaldehyde stress, and further demonstrated the importance this protein plays at both transcriptional and translational levels in this model methylotroph.

## INTRODUCTION

Stressors in biological systems can have multiple, simultaneous effects on cells. They can have single or numerous molecular targets, each of which may affect different cellular processes, and ultimately the consequences can radiate throughout metabolic and regulatory networks to the whole cell. Antibiotics, for example, inhibit particular proteins or RNAs, which themselves are involved in processes including the synthesis of genomes, proteins, or cell walls [1]. Ultimately, this leads to a slowing or cessation of growth. On the other hand, stressors such as heat, osmotic pressure, radioactive radiation, pH change, exposure to toxic metals or aldehydes do not have specific targets; all of these examples cause proteins to misfold extensively [2–6]. Although there are still layers to the response – the proteins that are misfolded, and the processes affected by them are impaired – both the direct and indirect dimensions of the cellular response are broad, with many proteins and processes affected simultaneously at both levels.

Formaldehyde is a potent toxin that naturally occurs in biological systems. It is typically generated as a byproduct of enzymatic reactions such as demethylation reactions and heme degradation or by non-enzymatic breakdown of metabolites such as glycine, tetrahydrofolate, and methanol. Formaldehyde notoriously reacts with DNA, yielding a variety of DNA modifications including intrastrand crosslinks and DNA-protein crosslinks [7–9]. However, its electrophilic nature also makes it highly reactive with free amines and thiols, and it likely reacts with myriad cellular molecules and disrupts other cellular processes [10,11]. To prevent the inevitable cellular damage that would result from elevated levels of this cytotoxin, many organisms are equipped with formaldehyde detoxification pathways that convert formaldehyde to more benign metabolites, such as formate, that can be further modified by cellular metabolism [12–14].

In bacteria, there are four major classes of formaldehyde detoxification pathways [12]. Thiol-dependent pathways are the most prevalent [15–19]. They involve the condensation of formaldehyde and the major cellular thiol, typically glutathione. In this manner, formaldehyde is ultimately oxidized to formate and the thiol compound is regenerated. Similarly, pterin-dependent pathways, which require tetrahydrofolate or tetrahydromethanopterin, use these one-carbon carriers to oxidize formaldehyde to formate or other biosynthetic intermediates [20,21]. The ribulose monophosphate-dependent pathway fixes formaldehyde to ribulose monophosphate and the resulting product, *D*-arabino-3-hexulose-6-phosphate, is converted to fructose-6-phosphate which can enter catabolic pathways such as glycolysis [22]. Lastly, whereas all of the above pathways involve attaching formaldehyde to a cofactor or phosphosugar for detoxification, some organisms also encode aldehyde dehydrogenases that can act directly upon formaldehyde, although evidence for their physiological role is sparse [23].

In methylotrophs, formaldehyde is central to cellular metabolism and not simply a nuisance byproduct or degradation product [24–26]. Methylotrophs can use reduced one-carbon compounds as sole sources of carbon and energy, and they almost universally convert these one-carbon units to formaldehyde. Thus, formaldehyde is an obligate central intermediate with the potential to become an internal stressor and a potentially lethal toxin in these organisms. In *Methlyorubrum extorquens*, the most extensively studied facultative methylotroph, the primary mechanism for formaldehyde detoxification is the formaldehyde oxidation pathway which is mediated by dephospho-tetrahydromethanopterin (dH_4_MPT) [24]. Therefore, during growth on methylotrophic substrates such as methanol, the carbon utilization pathway and formaldehyde detoxification pathway are one and the same. Previous work showed that when mutants that cannot synthesize dH_4_MPT or lack enzymes of the dH_4_MPT-dependent pathway were actively growing on succinate and then suddenly exposed to methanol they experienced a temporary halt in growth [25]. This methanol-sensitivity was dependent on methanol dehydrogenase, suggesting that the inability to utilize formaldehyde led to toxicity, likely from a rapid increase in internal formaldehyde [25]. When *M. extorquens* is growing on methanol, intracellular formaldehyde concentrations are relatively high, approximately ~1 mM [27]. In spite of the central role of formaldehyde in methylotrophic metabolism, little is known about how methylotrophs evade or reverse formaldehyde-inflicted damage when the metabolic network transiently becomes imbalanced, as might occur during nutrient shifts or under dH_4_MPT limitation.

In *M. extorquens* PA1, the functions contributing to formaldehyde toxicity were probed through experimental evolution for growth on formaldehyde. Evolution to grow on 20 mM formaldehyde [28] commonly resulted in loss-of-function mutations in a gene that has since been renamed *efgA* (for enhanced formaldehyde growth). Further experimentation showed that EfgA directly binds formaldehyde and interacts with the cell’s translational machinery to rapidly arrest translation in the presence of excess formaldehyde. The working model is that EfgA senses excess formaldehyde and inhibits translation to prevent cellular damage (Figure 1). Halting translation may mitigate a specific type of damage the cell is experiencing, namely proteotoxicity. However, it is also possible that this novel regulatory mechanism is used because it is a rapid way to shut down cellular activity to prevent widespread damage and maintain viability. To further characterize the protective role of EfgA, we sought to investigate the global response of *M. extorquens* to formaldehyde by eliminating the primary mechanism that *M. extorquens* uses to prevent formaldehyde stress and investigating the cellular response to formaldehyde in its absence.

**Figure 1.**
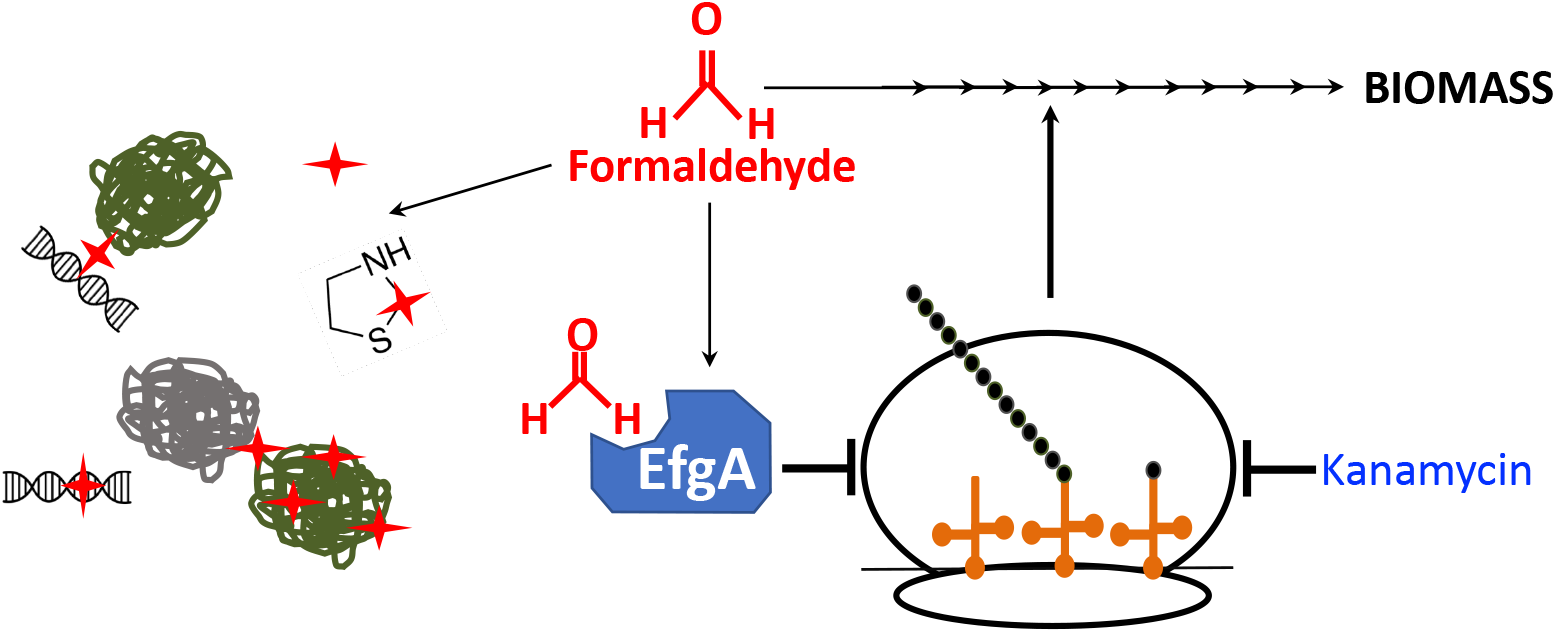
Fate of formaldehyde in *M. extorquens*. In *M. extorquens*, formaldehyde is oxidized and used as a source of carbon and energy for biomass production. When levels of formaldehyde are excessive, free formaldehyde can form adducts on and/or crosslink a number of molecules, including proteins and nucleic acids, via amine and thiol groups. EfgA is a formaldehyde-sensing protein that, when bound to formaldehyde, halts translation.

Here, we investigate growth and gene expression in *M. extorquens* PA1 in response to formaldehyde stress. Our outstanding questions at the onset of this work were: What are the consequences of formaldehyde stress with and without EfgA, the cell’s primary avenue to sense and respond to this stress? Are cells metabolically active during the formaldehyde-induced stasis and how do they recover from formaldehyde stress and resume growth? Furthermore, how does EfgA-formaldehyde-mediated translational inhibition compare to other mechanisms of translational inhibition, such as exposure to particular antibiotics? Thus, we sought to disentangle the cellular responses that are specific to formaldehyde, mediated by EfgA, from those generalizable to translation inhibition. To achieve this, we compared the response to formaldehyde in wild type and a Δ*efgA* mutant over an extensive timecourse. In parallel, we compared the response of each strain to the translational inhibitor kanamycin. We found that in actively growing cells, formaldehyde stress rapidly induces quiescence, a metabolically-active non-replicating state. The transition to a quiescent state requires EfgA, which mediates 70% of the observed global response to formaldehyde, which is subsequently reversed when levels of formaldehyde present decrease. In the absence of EfgA, *M. extorquens* has a delayed and muted response to formaldehyde and exhibits signs of proteotoxic and genotoxic stress. These findings were in stark contrast to the cellular response to kanamycin where the timescale of the transcriptional response was on the order of hours and lethality was on the order of minutes; however, both mechanisms of translation inhibition also had overlapping features.

## METHODS

### Bacterial strains, media, and chemicals

Strains used in this study are derivatives of *Methylorubrum extorquens* PA1 [29, 30]. The ‘wild-type’ strain (CM2730) lacks genes for cellulose synthesis (Δ*celABC*) to prevent biofilm formation and enable reproducible growth [31]. The Δ*efgA* mutant (CM3745) additionally has a markerless deletion at the *Mext_4158* locus which encodes EfgA [28]. All experiments were performed with *Methylobacterium* PIPES (MP) medium [31] with 3.5 or 15 mM succinate as a carbon source. Bacto Agar (15 g/L, BD Diagnostics) was added for solid medium. Formaldehyde stock solutions (1 M) were prepared by boiling a mixture of 0.3 g paraformaldehyde and 10 mL of 0.05 N NaOH in a sealed tube for 20 min. When present in media, these compounds were added to the following final concentrations: formaldehyde (5 mM), and kanamycin (50 μg/mL). All chemicals were purchased from Sigma-Aldrich unless otherwise stated.

### Growth quantitation

Strains were inoculated into 2 mL MP medium with 3.5 mM succinate in biological triplicate from individual colonies. Cultures were grown at 30 °C with shaking for 24 hr and then subcultured (1/64) into 2 L of MP medium with 15 mM succinate. Cell density was quantified by monitoring optical density (OD) at 600 nm (OD_600_). Specific growth rates (μ; per hour) were determined as follows: μ = ln[(*X*/*X0*)/*T*], where *X* is OD_600_ during exponential growth, *X*0 is OD_600_ at time zero, and *T* is time (in hours). To monitor cell viability, cells from a 100 μL aliquot of culture were harvested via centrifugation, discarding supernatant and resuspending cell pellets in MP medium (with no carbon source). Cell suspensions were then serially diluted (1/10 dilutions, 200 μL total volume) in 96 well polystyrene plates with MP medium (no carbon), 10 μL aliquots of each dilution were spotted to MP medium plates (15 mM succinate) in technical triplicate. Plates were incubated at 30 °C and colony forming units were quantified on days 3 and 6 of incubation. Technical triplicates were averaged for each biological replicate.

### Formaldehyde quantification

Formaldehyde concentrations in the culture media were measured as previously described [32]. Briefly, a 100 μL aliquot of culture was centrifuged to separate cells and supernatant. In technical triplicate, 10 μL of the supernatant or 100 μL of 0.1X supernatant (diluted with MP medium, no carbon) was combined with 190 or 100 μL Nash reagent B, respectively, in 96 well polystyrene plates. Plates were incubated (60 °C, 10 min), cooled to room temp (5 min) and absorbance was read at 432 nm on a Wallac 1420 VICTOR Multilabel reader (Perkin Elmer). Formaldehyde standards were prepared daily from 1 M formaldehyde stock solutions and a standard curve was generated in parallel with samples at each measurement (timepoint).

### Transcriptomic analysis

#### Cell treatment

From individual colonies, wild type (CM2730) and the Δ*efgA* mutant (CM3745) were inoculated into 2 mL MP medium (3.5 mM succinate) and grown at 30 °C with shaking for 32 hr. Cells were subcultured (1/64) into fresh MP media twice: first into 35 mL (3.5 mM succinate, 24 hr incubation) and then into 2 L (15 mM succinate). When OD_600_ of 2 L cultures = 0.2 (~8 hr), 100 mL of each culture was used to harvest “pretreatment” cells for transcriptomic analysis and remaining cultures were divided into 100 mL aliquots in 250 mL flasks and returned to the incubator for 30 min. All flasks were quickly removed from incubator and treated with i) 500 μL of 1 M formaldehyde (5 WT flasks, 4 Δ*efgA* mutant flasks), ii) 100 μL of 50 mg/mL kanamycin (4 flasks of each strain), or iii) left untreated (4 flasks of each strain). Flasks were sealed with foil and parafilm and returned to the shaker. Time of treatment is designated as t = 0, pretreatment therefore corresponds to −45 min. This growth regimen was executed on 6 consecutive days to obtain all treated samples in biological triplicate and yielded 6 biological replicates of “pretreatment” samples. Cells of each strain and treatment were harvested at 5, 20, 40, 60, 90, 180, 360, and 540 min.; formaldehyde-treated wild-type cells were additionally harvested at 720 and 1080 min.

#### Harvesting cells

At each time point a 100 mL culture was divided into 2-50 mL conical vials (on ice). Cells were pelleted by centrifugation (3220 x *g* 5 min, Eppendorf 5810R fitted with rotor A-4-81), pellets were recombined with 5 mL pre-chilled MP medium (no carbon), and evenly divided into 5-2 mL cryovials (on ice). Cells were pelleted by a second round of centrifugation. After discarding the supernatant the cryovials were resealed and submerged in liquid nitrogen. Cell pellets were stored at −80 °C.

#### RNA sequencing

RNA was purified using the RNAsnap method [33]. First, cell pellets were resuspended in 500 μl of RNA extraction buffer (18 mM EDTA, 0.025% w/v SDS, 1% v/v 2-mercaptoethanol, 95% v/v formamide) by vortexing. Following incubation at 95°C for 7 min, cell debris was pelleted by centrifugation at 16000 × *g* for 5 min at room temperature. The resulting supernatant was purified using the Clean & Concentrator kit incorporating the on-column DNase digestion step (Zymo Research). Yields were determined using a Qubit 2.0 fluorometer with the RNA BR assay kit (ThermoFisher). Then, 1.5 μg of purified RNA for each sample was processed using the Ribozero rRNA Removal Kit (Gram-negative bacteria) (Illumina) according to the manufacturer’s instructions except that reaction volumes were reduced by 50%. Depleted RNA samples were ethanol precipitated.

RNA-Seq library preparation from rRNA-depleted samples began by fragmenting them with the NEBNext Magnesium RNA Fragmentation Module (New England Biolabs) for 110 s followed by ethanol precipitation. Fragmented RNA was treated with 10 U of T4 polynucleotide kinase (NEB) in 20 μl reactions containing 1 mM ATP. After another ethanol precipitation, the NEBNext small RNA Library Prep Set for Illumina (Multiplex Compatible) (New England Biolabs) was used to prepare samples according to recommended protocol, except all reaction volumes were reduced 50% and the SR RT primer concentration was reduced by an additional 50% in the RT primer hybridization step. After performing 15 cycles of PCR, samples were ethanol precipitated, and the DNA concentration was determined via the Qubit HS dsDNA assay kit (Thermofisher). For each sample, 200 ng of DNA was run on a 4% EX E-gel (ThermoFisher). DNA with a size of at least ~200 bases was excised and purified using a gel DNA recovery kit (Zymo Research) with elution in 50 μl of nuclease-free water. These DNA libraries were subjected to paired-end 150-base sequencing on an Illumina HiSeq 4000 or single-end 75-bp sequencing on an Illumina NextSeq 500 by the Genome Sequencing and Analysis Facility at the University of Texas at Austin.

#### Data analysis

##### Sequence alignment and determination of counts per gene

Illumina reads were trimmed of adaptors using Trimmomatic v0.38 [34]. We used the paired-end trimming mode, if applicable, and settings that removed reads trimmed to fewer than 30 bases. Remaining read pairs and singleton reads were mapped to the *M. extorquens* PA1 genome (GenBank: NC_010172.1) [29] using Bowtie2 v4.8.2 [35] with default settings, except we used the option to report only one alignment per read. The resulting SAM files were processed using HTSeq v0.11.4 [36] to count read pairs or singleton reads that unambiguously overlapped each gene. Scripts with the commands for running this analysis are available in GitHub (https://github.com/barricklab/brnaseq).

##### Normalization of the data and Principal Component Analysis (PCA)

All data manipulation and statistical analysis was done in R version 3.6. The matrix of raw count data was converted to normalized counts using DESeq2 [37]. For PCA, all the treatments were normalized against each other. For plotting, the plotPCA function in DESeq2 was used to generate the PCA dataset and the result was used in ggplot2 package to make the plot.

##### Heatmaps and Venn Diagrams

For heatmap plots, each treatment was normalized against WT pre-treatment (the control pre-treatment at 45 minutes before stressor was added to any treatment). To reduce the number of false-positive genes across the ~5000 genes in the genome, we used a very conservative criterion: subsets of genes with False Discovery Rate (FDR) adjusted p-values less than 0.001 were selected. Heatmaps were generated using the heatmap.2 function in the gplots package [38]. To do pairwise comparisons between two treatments (or two categories of treatments) we used the heatmap data to plot the Venn diagrams. Based on the sign of fold change, genes were divided into up or down-regulated categories. The VennDiagram package [39] was used to generate the data needed for plotting.

##### Analysis of significance in candidate genes

For the modest number of candidate genes examined, the Wald test was used to calculate the significance of the change in mRNA levels (Log2FC). Genes with FDR adjusted p-values less than 0.001 were classified as significantly changed genes. Furthermore, for the formaldehyde metabolism genes, we used a one-tailed test because we had a specific direction for our *a priori* hypothesis. Genes with both positive fold changes and positive Wald-statistics were classified as up-regulated, and those with negative fold changes and negative Wald-statistics were classified as down-regulated.

Raw Illumina read data and processed files of read counts per gene and normalized expression levels per gene have been deposited in the NCBI GEO database (accession GSE163955).

### Gene enrichment

The gene sets of differentially expressed genes, were examined for the enrichment of genes with particular functions using **D**atabase for **A**nnotation, **V**isualization and **I**ntegrated **D**iscovery (**DAVID**) version 6.7 [40, 41] and Comparative GO (comparativego.com) [42, 43]. Both methods produced comparable groups of gene sets (Tables S1, S2). Figures were generated using GraphPad Prism.

## RESULTS

### Formaldehyde stress induces a quiescent state

Previous work demonstrated that growth and translation were inhibited in an EfgA-dependent manner when exponentially growing *M. extorquens* was treated with formaldehyde [28]. Here we examined the effects of formaldehyde treatment on the growth dynamics of wild type *M. extorquens* and a Δ*efgA* mutant by monitoring cell density, viability, and formaldehyde concentrations in the growth medium in cultures over an extended timecourse. Strains were grown in minimal medium containing succinate. Under these non-stressful conditions, formaldehyde is not generated from carbon utilization in any appreciable amount. Once cultures reached an early exponential phase of growth (OD_600_ ~ 0.2), they were divided and left untreated or were treated with 5 mM formaldehyde. As anticipated, when wild type and Δ*efgA* mutant cells were untreated, the growth of both strains was identical and continued to be exponential, as indicated by both cell density and viability (Figure 2A, 2B). This suggests that, in the absence of formaldehyde, the absence of EfgA does not impact growth or viability. By contrast, when cells were treated with 5 mM formaldehyde, the effect of EfgA was apparent. The formaldehyde-treated wild-type cells were immediately growth arrested while the formaldehyde-treated Δ*efgA* mutant grew nearly as well as the no-stressor control, only experiencing a 14% reduction in growth rate relative to the wild type (! = 0.2140 versus 0.1840, p-value = 0.007). Growth arrest in wild-type cells lasted for at least 360 min, as was evident by unchanged OD and cell viability over this time period. Importantly, during that time cells did not show any evidence of loss of viability.

**Figure 2.**
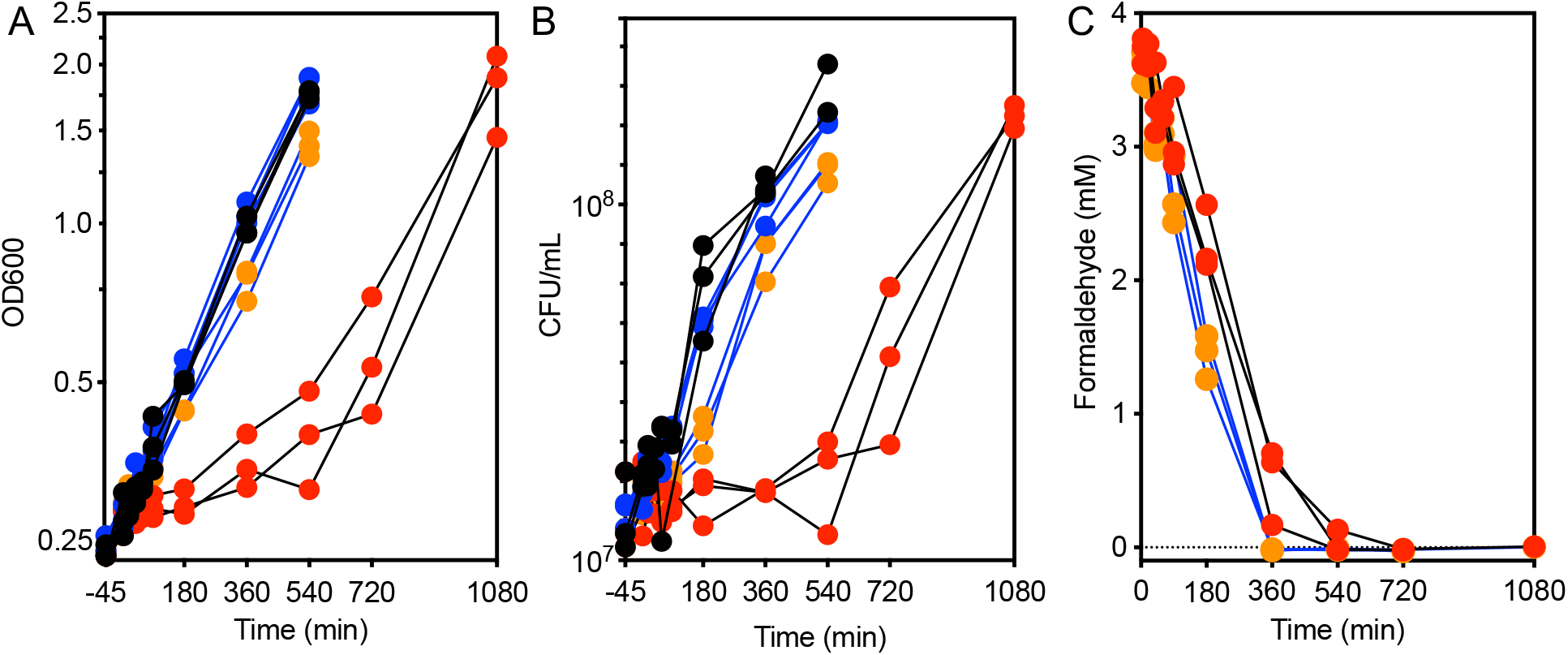
Formaldehyde induces EfgA-dependent growth arrest. The growth response of WT (black lines) and Δ*efgA* mutant (blue lines) was quantified in minimal succinate medium with no addition (black/blue symbols) and with formaldehyde (red/orange symbols). Growth was assayed by A) optical density (OD_600_) and B) cell viability (CFU/mL). Formaldehyde concentrations in the culture supernatant were measured in formaldehyde-treated samples (C).

Formaldehyde concentrations in the growth medium were monitored by a colorimetric (Nash) assay. The resulting data showed that formaldehyde was gradually depleted in both the wild type and Δ*efgA* mutant cultures. Despite being growth arrested, wild-type cells depleted formaldehyde from the growth medium nearly as quickly as the Δ*efgA* mutant (WT = 0.54 ± 0.03 mM/h, Δ*efgA* = 0.62 ± 0.02 mM/h), suggesting they were in a quiescent, metabolically active state [44]. These data suggest that upon formaldehyde stress, *M. extorquens* cells can detoxify their environment and passively wait out this exogenous stressor.

### Cells resume growth when the formaldehyde stress is reduced

After a prolonged period of growth arrest, wild-type cells resumed growth between 360 and 540 min (6 and 9 hr) post-formaldehyde treatment. Within that timeframe, formaldehyde concentrations in the growth medium had plummeted to non-inhibitory concentrations (<1 mM) (Figure 2C). By contrast, when the Δ*efgA* mutant was challenged with formaldehyde, it continued to grow; however, when compared to untreated samples, the cell density and viability at the final timepoint (540 min) were compromised (78% and 74% of that of untreated samples, respectively). These data suggested that in the presence of EfgA, formaldehyde is a titratable signal whose absence permits cells to turn growth back “on” after formaldehyde-induced stasis. Furthermore, the suboptimal growth of the Δ*efgA* mutant upon formaldehyde treatment suggests that in the absence of EfgA, while cells may still grow, they experience formaldehyde stress/damage as a result of not arresting growth.

### Initial experiments to identify timescale of response for further transcriptomic analyses

To evaluate the global response to formaldehyde, we examined the transcriptome of the wild type and the Δ*efgA* mutant during this timecourse and compared it with that of the global response to kanamycin, an aminoglycoside antibiotic that inhibits translation by binding to the ribosome. Initially, we performed RNA-sequencing on individual replicates that spanned the breadth of experimental variables (genotype, treatment, time) and identified samples where the earliest and most intense responses were observed. In this single replicate analysis, we found that exponential-phase wild-type samples, taken after formaldehyde depletion, had few differences relative to the pretreatment samples (Figure S1). Thus, it seemed that there was no large physiological event that mediated growth resumption but that it simply reflected a reversal of the formaldehyde-EfgA inhibition effect. Unexpectedly, with kanamycin treatment, we saw minimal differences at the 40 min timepoint with the single replicate in comparison to the formaldehyde treatment at the same timepoint. As a result, we limited our analyses of formaldehyde-treated samples to those leading up to the 360 min timepoint and for the kanamycin-treated samples to the ≥40 min post-treatment samples. For the subset of samples we wanted to investigate further, we performed RNA-sequencing in biological triplicate.

### Overall pattern of gene expression changes revealed via Principal Component Analysis

To assess the variance among samples and overall trends, we performed principal component analysis (PCA) of all of the normalized, log-transformed triplicate data. Here we found that our biological replicates were consistently well-clustered (Figure 3). We also found large-scale patterns that showed that pretreatment and untreated samples clustered with each other independent of genotype or time, as did some treated samples from later timepoints (Δ*efgA* at 180, 360 minutes). In the earlier formaldehyde-treated samples, both genotypes deviated significantly from untreated samples and from each other. In each instance, after reaching their maximal distance from the untreated samples, they successively clustered closer to the untreated samples. The PCA analysis also showed that upon kanamycin treatment (discussed in detail in a later Results section) and formaldehyde treatment, the PC1 appears to have captured the strength of the shared response seen for both kanamycin and formaldehyde as stressors while PC2 appears to distinguish the two stressors from each other.

**Figure 3.**
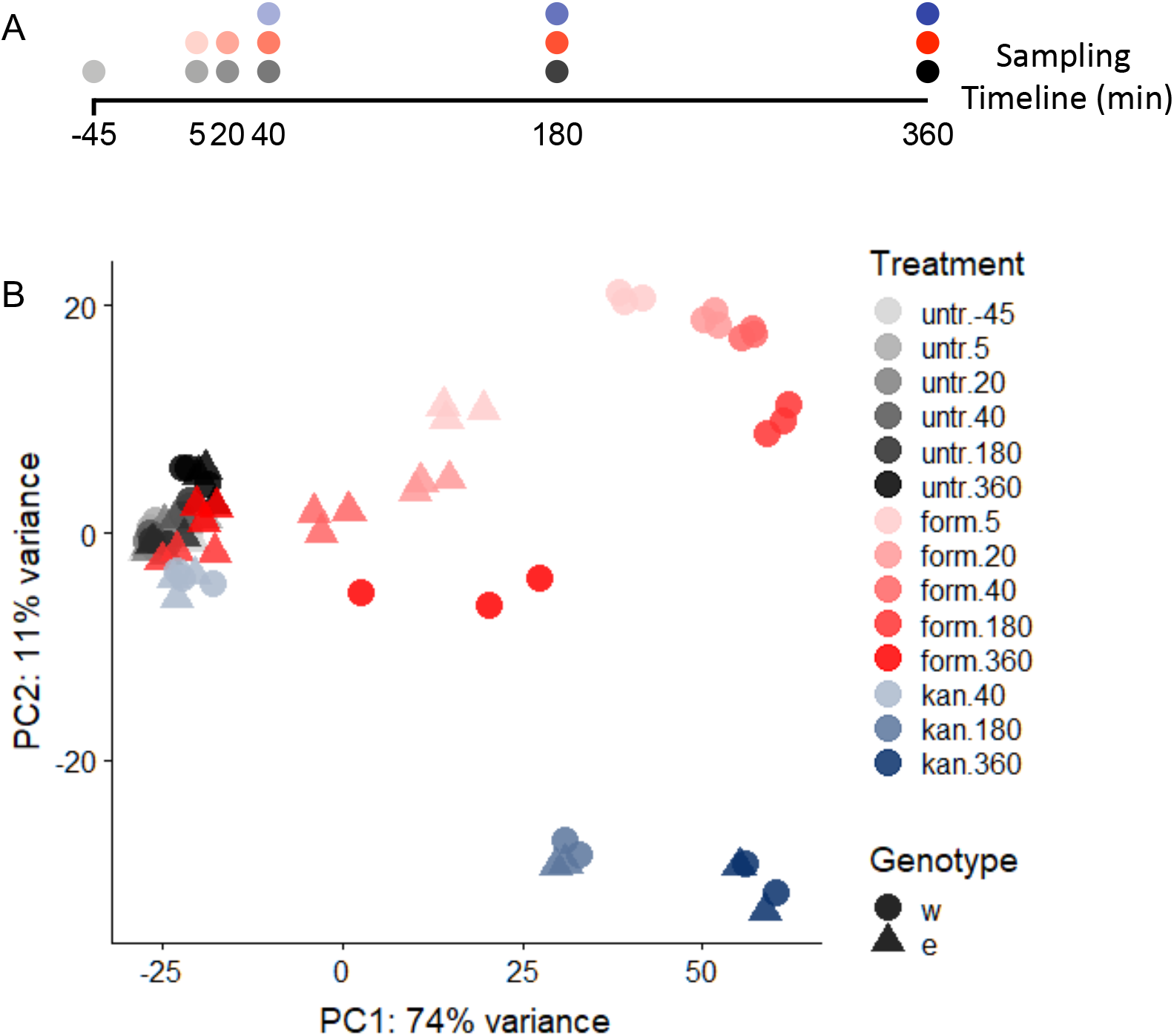
PCA of WT and Δ*efgA* mutant transcriptomes. Timecourse RNA-Seq was performed on WT (circles) and Δ*efgA* mutant (triangles) of *M. extorquens* grown in succinate minimal medium. During early exponential growth cultures were left untreated (untreat, gray gradient), or treated with 5mM formaldehyde (form, pink gradient) or 50ug/mL kanamycin (kan, blue gradient). Principal Component Analysis (PCA) was performed to assess the variance in the data. Each symbol represents one experimental replicate; the colors of each gradient darken over exposure time (-45 (pretreatment) to 360 min).

### *efgA* genotype does not impact the transcriptome during growth upon succinate in the absence of formaldehyde

Although growth of WT and the Δ*efgA* mutant in succinate when untreated was similar to each other and consistent through time, we first sought to determine whether there were underlying transcriptional differences between the genotypes in the absence of stressor or as either grew throughout the experiment. To identify differentially expressed genes, we imposed a two-fold change cutoff (Log2FC > 1.0) with an adjusted p-value less than 0.001. Prior to evaluating the effects of each treatment, we performed a pair-wise comparison of the log-transformed counts of wild-type untreated sample during exponential growth (5, 20, 40, 180 min) versus pretreatment and found that 22 genes were differentially expressed in at least one timepoint (Table S3). We performed the same comparison with the Δ*efgA* mutant and found that 21 of the 22 genes identified in the wild-type samples were also differentially expressed and that the directions of these changes were conserved. Additionally, we found only six other genes were ever differentially expressed in the Δ*efgA* mutant (Table S3, bolded). Therefore, only 0.4% of the genes in wild type and 0.5% of the genes in the Δ*efgA* mutant showed a significant change at any time from 5 minutes to 180 minutes. These data suggested that during growth on a non-methylotrophic growth substrate (i.e., in the absence of any formaldehyde stress), the transcriptome of the wild type and Δ*efgA* mutant are nearly identical and the presence of EfgA has a minimal impact on global gene expression. By 360 min, there were 3.9% up-regulated and 1.3% down-regulated genes in WT, and 5.3% up-regulated and 3.8% down-regulated genes in the Δ*efgA* mutant. Of these genes, 48.7% of up-regulated genes and 49.2% of down-regulated genes were in common between the two genotypes at 360 minutes. These findings allowed us to simplify the analyses with stressors described below by simply using the wild-type pretreatment sample as the reference point for all subsequent analyses (i.e., the control).

### The global response to formaldehyde is extensive and predominantly mediated by rapid action of EfgA

In wild type, by the earliest timepoint after formaldehyde treatment (5 min), cells were already differentially expressing 1,783 genes, which is 37% of the 4,829 protein-coding genes in *M. extorquens*. Of these, 935 genes were upregulated and 848 were downregulated (Figures 4, 5, S2). Over the next three timepoints (20, 40, 180 min) the response intensified both in the number of genes differentially expressed (2,246) and the magnitude of the expression changes. A key feature in the response to formaldehyde was that the response in each timepoint was a superset of the previous timepoint. The dramatic shift in the transcriptome began to subside by the 360 min time point (down to 18% of genes being differentially regulated, from a high of 47% at 180 min); this coincides with the previously described timepoint where formaldehyde was nearly cleared from the supernatant (<1 mM) and growth had resumed.

**Figure 4.**
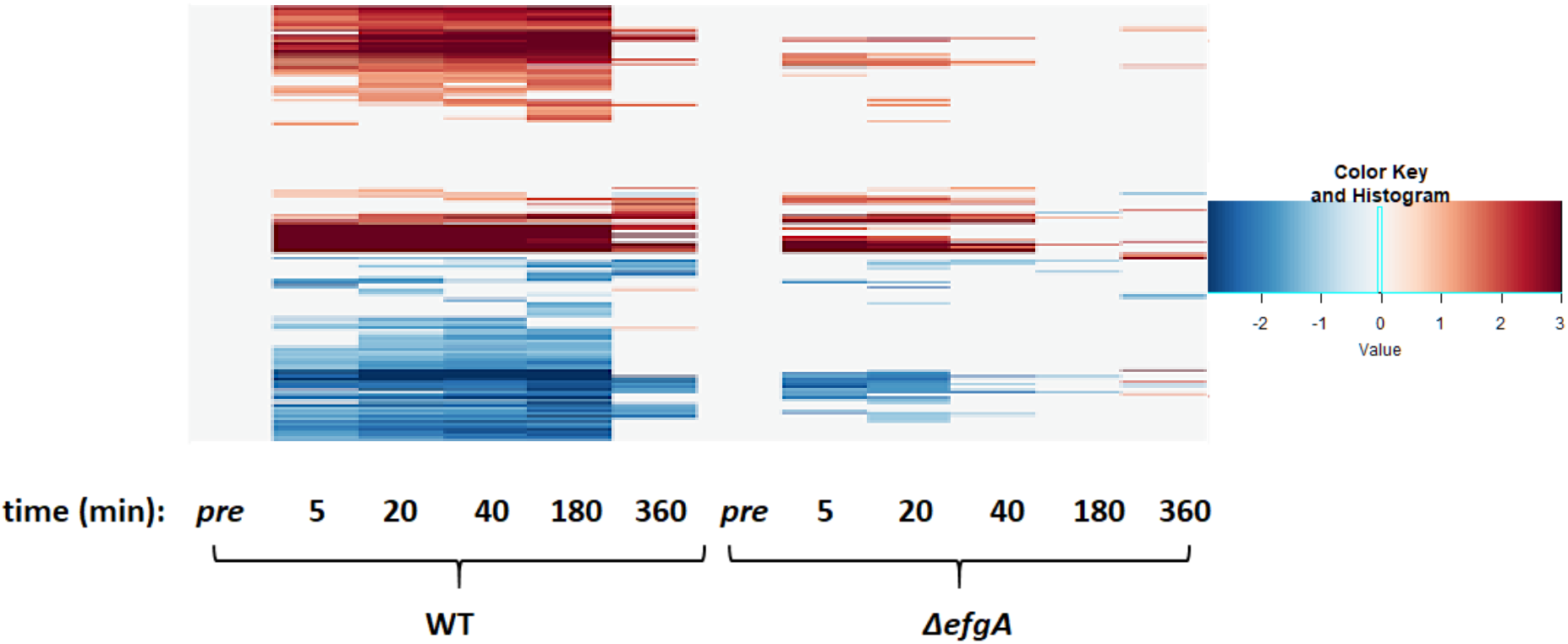
Temporal heatmap of WT and Δ*efgA* mutant response to treatment with formaldehyde. Genes differentially expressed (|log2-fold change| >1) were identified by dividing Log2FC of sample/Log2FC of pretreatment sample (of the respective genotype). The up or down directionality of expression change is indicated by the red or blue color gradient, respectively.

**Figure 5.**
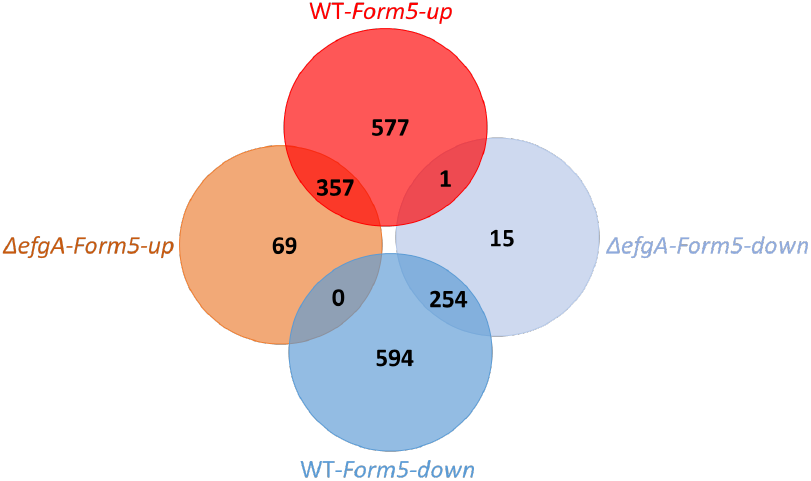
Venn diagram of genes differentially expressed in WT and Δ*efgA* mutant upon formaldehyde treatment. Genes differentially expressed (|log2-fold change| >1) were identified by dividing Log2FC of sample/Log2FC of pretreatment sample (of the respective genotype). Venn diagrams for later timepoints are depicted in Figure S2.

In the Δ*efgA* mutant, the response to formaldehyde was less extreme than that of wild type and subsided faster. The mutant differentially expressed 696 genes (~14% of protein-coding genes in *M. extorquens*). This represented ~39% of the number of genes differentially expressed in wild type, though some expression changes were unique to the Δ*efgA* mutant. Additionally, in the Δ*efgA* mutant expression generally had more modest fold changes than wild type (Figures 4, 5, S2). In the Δ*efgA* mutant, the number of differentially regulated genes peaked at 20 min, began to subside by 40 min, and was comparable to pretreatment samples by 180 min. In terms of intensity, the Δ*efgA* response got weaker from 5 minutes to 180 minutes, and the sets of genes that were differentially expressed at each time point were largely subsets of the ones from the previous timepoint. Collectively, these data indicate that the rapid response to formaldehyde is largely dependent upon EfgA. The EfgA-dependent response to formaldehyde accounts for 63% of the total cellular response.

### Loss of viability precedes the transcriptomic response to kanamycin

Given that EfgA appears to act via inhibiting translation in response to elevated internal formaldehyde, we sought to compare this response to the classical translation-inhibiting antibiotic, kanamycin, whose mechanism of action is to bind the 30S subunit of the ribosome and inhibit translation. Kanamycin-treated cells were derived from the same untreated wild type and Δ*efgA* mutant cultures described earlier that were partitioned for treatment. Upon kanamycin treatment, both genotypes exhibited comparable viability and transcriptomic responses.

Monitoring growth of kanamycin treated cells by OD showed that the ODs of both genotypes continued to increase post-treatment and coincided with that of the untreated samples for at least 180 min and then plateaued (Figure 6A). Cell viability assays, however, showed that by 40 min, the viability of both genotypes had decreased by an order of magnitude and continued to fall rapidly (Figure 6B). These seemingly conflicting growth data are not uncommon as each of these approaches measure different features of growth. In light of the viability measurements, changes in OD cannot be attributed to proliferation. The increase in absorbance suggests that cells are increasing in size, a phenomenon that has been associated with stress and unchecked translation [45–47]. By the time the ODs plateaued (180 min), viability had already decreased by four orders of magnitude.

**Figure 6.**
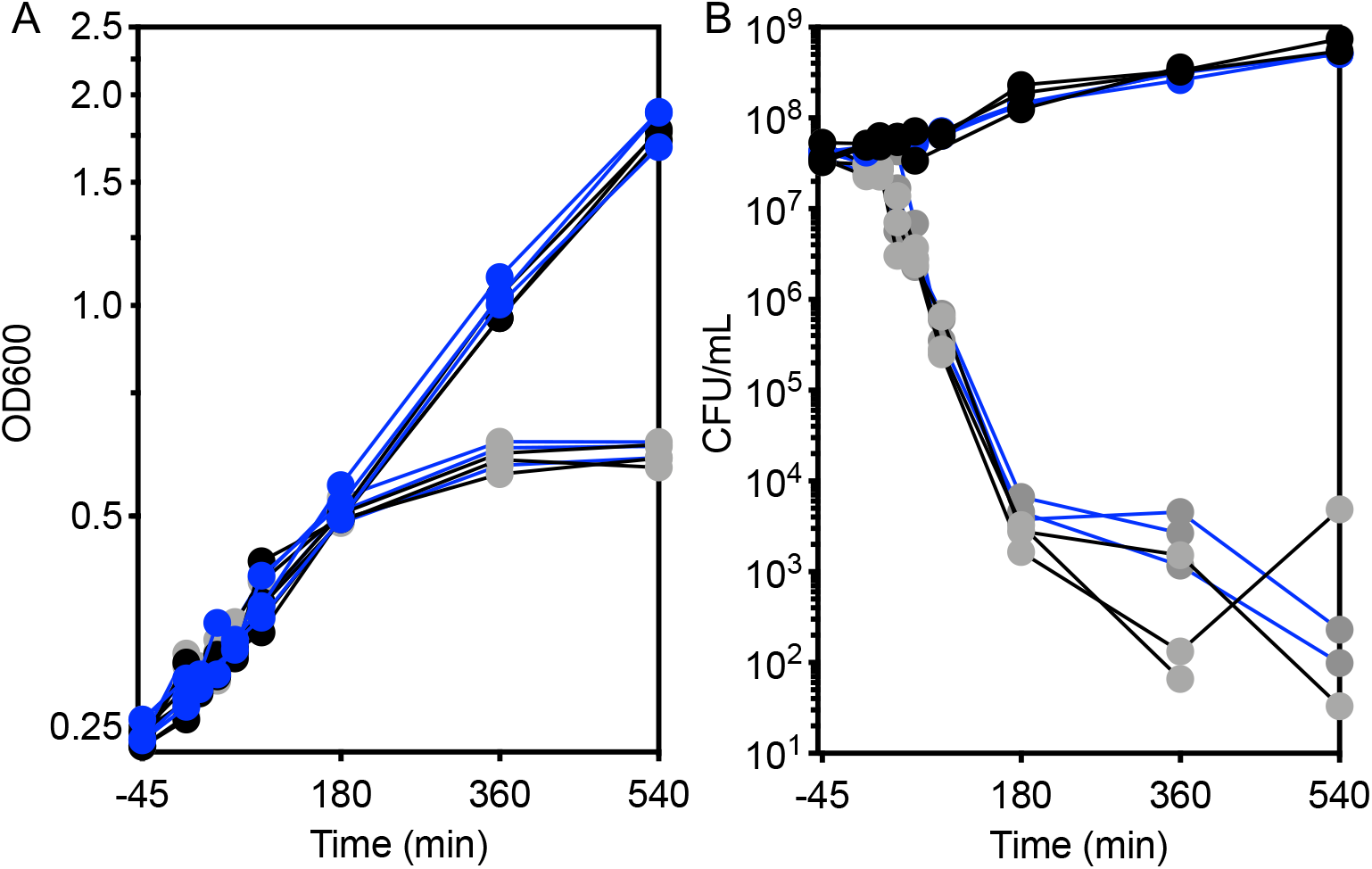
Kanamycin treatment results in early loss of viability. The growth response of WT (black) and Δ*efgA* mutant (blue) was quantified in minimal succinate medium with no addition (black/blue symbols) and with kanamycin (gray symbols). Growth was assayed by A) optical density (OD_600_) and B) cell viability (CFU/mL).

For transcriptomic analysis we used the wild-type untreated samples as the control for pairwise comparisons. Evaluation of the transcriptomes of the biological triplicates at 40, 180, and 360 min confirmed a near lack of a response at the 40 min timepoint with only 9 genes being differentially expressed. By 180 and 360 minutes, major changes in expression were observed (Figure S3). In WT, 30.1% and 40.6% of genes were differentially expressed at 180 minutes and 360 minutes, respectively. Similarly, in the Δ*efgA* strain 27.5% and 37.5% of genes showed significant changes in expression at 180 minutes and 360 minutes, respectively. For both strains, ~90% of the response at 180 minutes timepoint was a subset of the response that was amplified by 360 minutes (Figure S3). These data showed that the timescale of the formaldehyde response versus the kanamycin response are quite different. Among the functional categories that were enriched for in kanamycin-specific upregulated genes, unsurprisingly, were several related to translation and the ribosome (Figure S4). By contrast, enriched downregulated functional categories frequently contained metabolic genes, such as those related to the TCA cycle, cellular respiration, and glycolysis. This suggests that, transcriptionally speaking, kanamycin elicits a response focused on reviving translation while simultaneously attenuating cellular metabolism.

### Response to kanamycin and formaldehyde involve shared pathways

To assess the similarities and differences in the responses between these different mechanisms of translational arrest, we chose to compare the timepoints for wild type with the strongest overall responses to each stress: formaldehyde at 5 minutes, and kanamycin at 360 minutes. Remarkably, 46.3% of the total up-regulated genes and 51.9% of down-regulated ones were in common between these two treatments (Figure 7). Since the outcomes of viability are so different, it is likely that many of these common genes represent the consequences of inhibited translation. If this is the case, the key differences in how EfgA interacts with formaldehyde to cause an immediate, but non-lethal pause of translation are likely in the unique genes, or those that experienced opposite directions of change between the two stressors. Despite the large number of overlapping genes, these fell into few enriched functional categories, particularly for genes that were upregulated. Interestingly, among shared downregulated genes, there was strong enrichment in categories related to flagellar motility, signal transduction, and electron transport (Figure S5).

**Figure 7.**
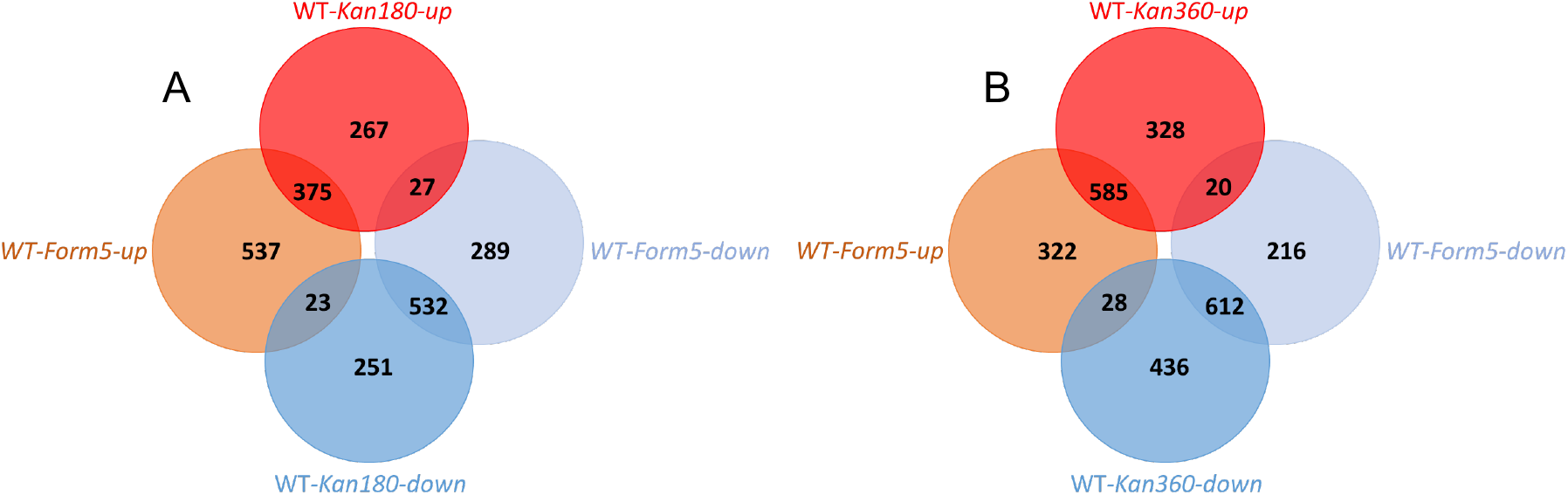
Response to kanamycin and formaldehyde has overlapping genes. Venn diagram of genes differentially expressed in WT upon formaldehyde treatment (5 min) or kanamycin treatment. Each diagram represents a different timepoint for the kanamycin treatment A) 180 min, B) 360 min. Genes differentially expressed (|log2-fold change| >1) were identified by dividing Log2FC of sample/Log2FC of pretreatment sample (of the respective genotype). The 40 min kanamycin timepoint was excluded as expression profiles had not yet changed significantly.

### Response of methylotrophy genes

Formaldehyde is an obligate metabolic intermediate of methanol utilization. As such, formaldehyde derived carbon and electrons are catabolized and ultimately assimilated into biomass. Thus, we expected that there would be changes in the expression of enzymes in the one-carbon metabolism pathways following formaldehyde treatment in our experiment. With succinate initially provided as the sole source of carbon and energy, there is the potential for formaldehyde utilization to occur simultaneously after it is introduced (i.e., co-consumption). Succinate and methanol are co-utilized in this way by *M. extorquens* [48]. Alternatively, formaldehyde can shock the system as a stressor, in which case formaldehyde oxidation might be primarily used as a detoxification mechanism. To query the cellular response with regard to carbon substrate utilization, we analyzed gene expression of methylotrophy genes. Methylotrophy can be broken down into several components: methanol oxidation, formaldehyde oxidation, formate oxidation, and formate assimilation through the tetrahydrofolate- (H_4_F)-dependent pathway and then the serine cycle (Figure 8). We assessed the cellular response to formaldehyde for each of these metabolic functions.

**Figure 8.**
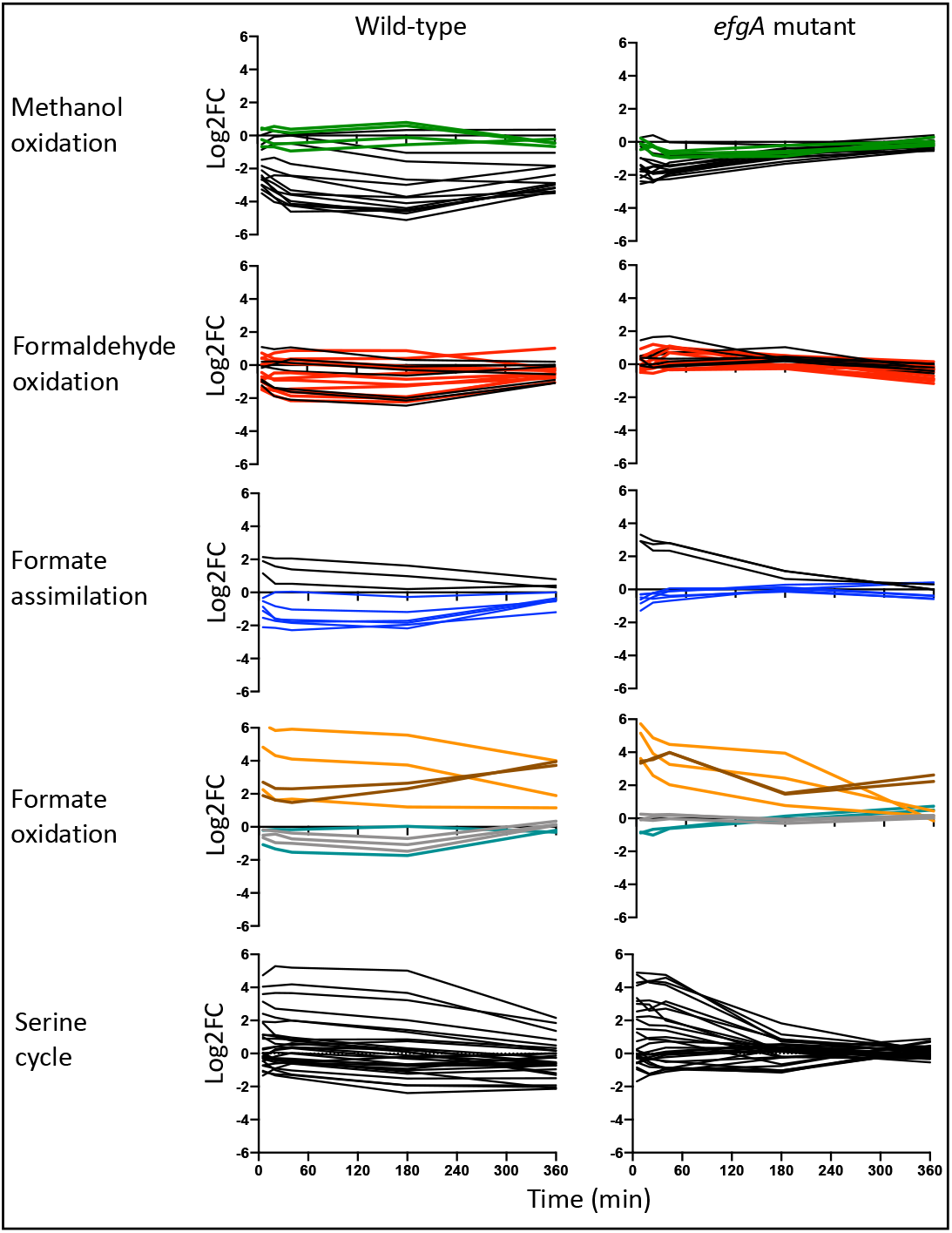
Temporal expression of genes of methanol utilization pathways. Temporal plot of the Log2FC of all genes involved in methanol utilization pathways in WT (left) and Δ*efgA* mutant (right). Each row represents genes involved in 1) methanol oxidation, 2) formaldehyde oxidation, 3) formate assimilation, 4) formate oxidation, and 5) serine cycle. Colors are used to distinguish genes involved in cofactor synthesis (PQQ synthesis, green; dH_4_MPT, red synthesis; blue, T_4_H synthesis) or different formate dehydrogenases (FDH1, teal; FDH2, orange; FDH3, gray; FDH4, brown).

#### Methanol oxidation

The *mxa* gene cluster encodes methanol dehydrogenase which is a PQQ-dependent enzyme that oxidizes methanol to formaldehyde during methanol utilization. When wild type is treated with formaldehyde, the *mxa* gene cluster is strongly downregulated with genes demonstrating up to a 34× decrease in expression. Expression remained downregulated up to 16× by the 360 min time point, when exogenous formaldehyde is near extinction and cells are resuming growth. The expression of the divergently transcribed methanol-regulated gene *mxaW* and PQQ biosynthetic genes were unchanged during the course of the experiment. In the *efgA* mutant, the *mxa* gene cluster is downregulated to a lesser extent (up to 5.5×).

#### Formaldehyde oxidation

Formaldehyde oxidation and detoxification involve the dH_4_MPT-dependent C_1_ transfer pathway. In wild type, expression of dH_4_MPT biosynthetic genes and those that encode dH_4_MPT pathway enzymes were not significantly different in untreated and formaldehyde-treated samples with the exception of *fhcABCD and OrfY*. The *fhc* genes, which encode the formyltransferase/hydrolase complex (Fhc) that releases formate, were downregulated up to 5.5×. In the *efgA* mutant, there were statistically significant but modest increases in expression of *fae*, *mtdB*, and *mch* at various time points but the largest contrast with wild type was seen in *fhcABCD* expression, which was unchanged. These data suggest that in response to formaldehyde, wild type may limit carbon flux by interfering with formaldehyde oxidation at the Fhc enzymatic step. A bottleneck imposed at the Fhc step could potentially restrict carbon flow by i) limiting the synthesis of downstream intermediates (formate, H_4_F pathway intermediates) and ii) leading to the accumulation of methenyl-dH_4_MPT, an intermediate in the dH_4_MPT pathway that regulates MtdA activity and prevents flux to assimilatory pathways via the Serine cycle [27].

#### Formate branchpoint

Formate is the main branchpoint in methylotrophic metabolism in *M. extorquens* where carbon flux can be partitioned to further generate energy by oxidation to CO_2_ or enter into a reductive pathway whereby formate carbon is transferred to tetrahydrofolate (H_4_F) carriers [26,49]. Upon introduction of formaldehyde, wild type increased expression of genes encoding formate assimilatory enzymes *ftfL*, *fch*, and *mtdA* but decreased the expression of H_4_F biosynthetic genes. Interestingly, assimilatory genes were also increased in the *efgA* mutant but the expression of the H_4_F biosynthetic genes was unchanged. As formaldehyde can spontaneously react with H_4_F to form methylene-H_4_F – the entry point of the serine cycle – at a slow rate, decreasing H_4_F biosynthesis may be a strategy for controlling flux to assimilation pathways or preventing accumulation of C_1_-H_4_F pool depletion in the presence of excess formaldehyde.

Overall expression of each of four formate dehydrogenases was comparable between genotypes. The expression of *fdh2* and *fdh4AB* were upregulated while that of *fdh1AB* was unchanged and *fdh3ABC* was modestly decreased (~2×).

#### Serine cycle

Because the wild-type strain reached a higher final cell density in the presence of formaldehyde, we concluded that formaldehyde carbon could be captured for biomass. However, as wild type was growth-arrested in the presence of formaldehyde for at least 6 hours we predicted that genes in the serine cycle pathway might be downregulated early on and increase later as cells resumed growth and used formaldehyde or formaldehyde-derived formate. However, in wild type, many serine cycle genes were strongly upregulated by the 5 min timepoint and remained elevated through the 360 min timepoint. The response in the *efgA* mutant was comparable to that of wild type during the early time points (5, 20, 40 min) but expression differences were widely gone by 180 min. Collectively, our data suggest that the regulation of transcription of serine cycle genes is not tightly linked to controlling carbon flux through the assimilatory pathways under these conditions.

Our data suggest that the *efgA* mutant simultaneously uses succinate and formaldehyde, which is consistent with previous work that demonstrates that when given the opportunity, *M. extorquens* will co-consume multi- and single-carbon growth substrates [48]. By contrast, wild type appears to delay consumption of and growth upon succinate until formaldehyde concentrations in the media have been drawn down to sub-inhibitory levels.

### Wild-type response to formaldehyde

To understand the wild-type transcriptional response to formaldehyde beyond methylotrophy genes, we identified functional classes of genes that were over-represented in our RNAseq analysis by gene enrichment using **D**atabase for **A**nnotation, **V**isualization and **I**ntegrated **D**iscovery (**DAVID**) [40, 41]. For genes differentially expressed at the 5 min timepoint, the Functional Annotation Clustering tool identified 61 clusters in the upregulated gene set, which included 793 genes, and 51 clusters in the downregulated gene set, which included 795 genes. DAVID could not annotate 10 and 53 genes in the upregulated and downregulated gene sets, respectively; thus they were excluded from clustering analysis. Each cluster was ranked by an enrichment score, the geometric mean (in log scale) of its component member's p-values; clusters that did not contain at least one member with p-value of <0.05 are not described here.

The clustering analysis revealed 14 enriched categories of upregulated genes and 21 categories of downregulated genes (Figure 9). The upregulated gene categories spanned diverse cellular processes from transcription to central carbon metabolism to metal binding. Due to their potential role in mitigating formaldehyde-induced damage, the DNA recombination/transposition and heat shock protein categories were notable (Figure 10). The DNA recombination category was composed of transposases (n=6), resolvases (n=3), integrases (n=2), and genes known to be involved in recombination pathways such as *recA* and *recO*. The heat shock category contains heat shock proteins (n=3), as well as the chaperones DnaK and ClpB, the latter of which can function as unfoldases and disassemble protein aggregates. The 21 downregulated categories included multiple aspects of signal transduction, including cyclic nucleotide metabolism and two-component regulatory systems as well cell wall biosynthesis and motility.

**Figure 9.**
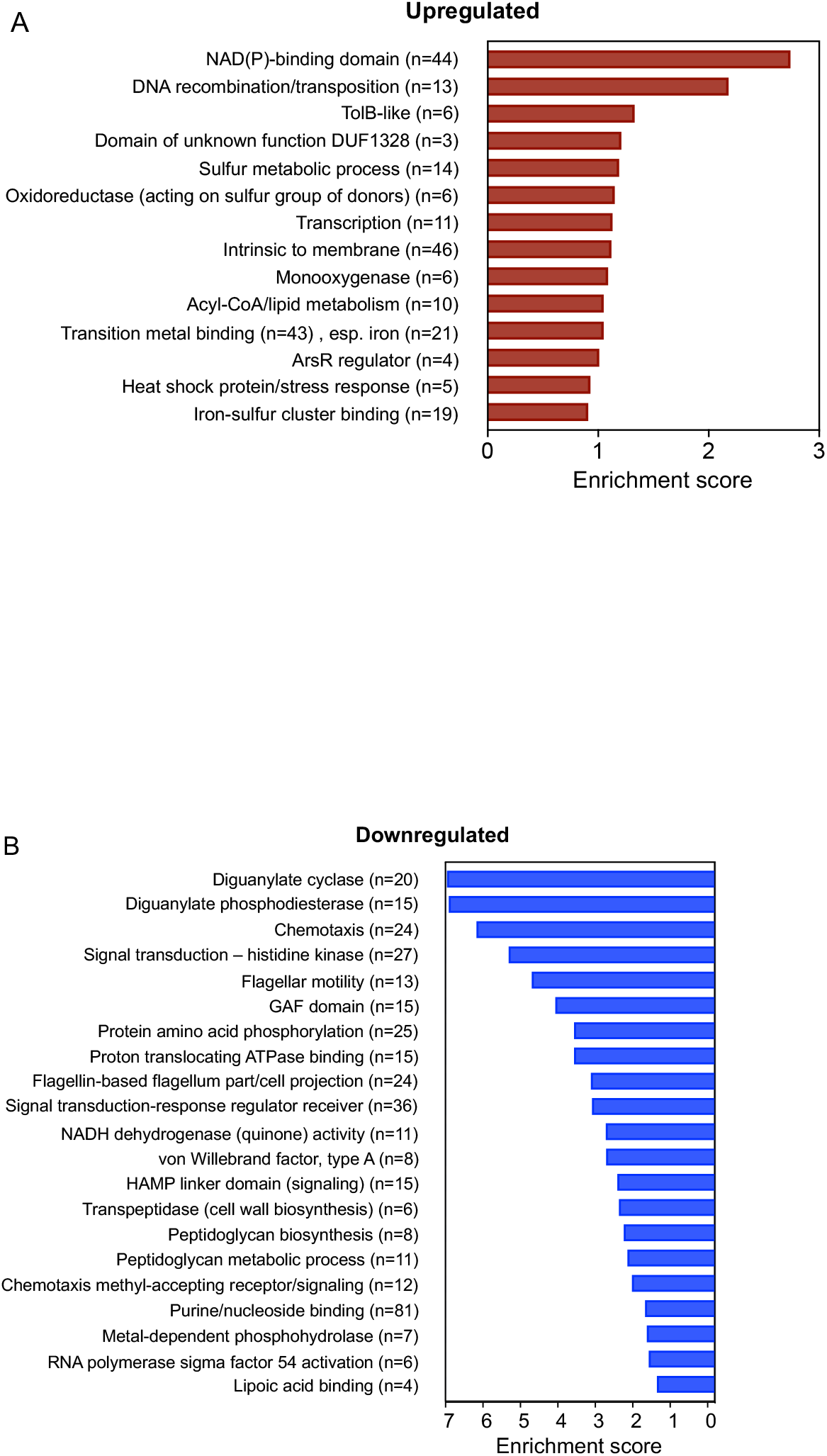
Gene ontology of the early formaldehyde response of WT. Summary of functional group response of WT at 5min (fastest response) and 20min (most intense response) was obtained using DAVID 6.7. Enrichment scores for each functional group is plotted for A) upregulated and B) downregulated genes; the number of genes in each functional group is indicated by ‘n’. All functional groups shown had p<0.05.

**Figure 10.**
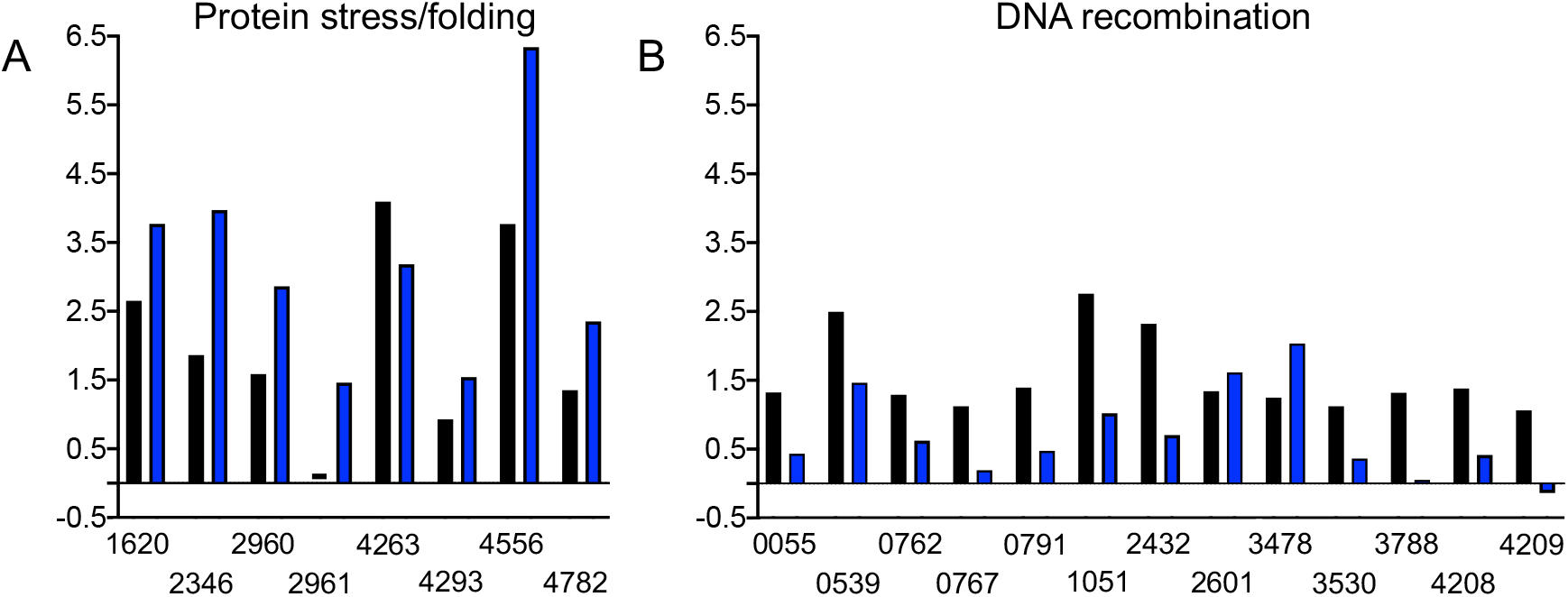
Genes involved in protein stress but not DNA recombination are upregulated in Δ*efgA* mutant in response to formaldehyde. Expression of genes involved in A) protein stress/folding and B) DNA recombination (as identified by gene ontology) are shown for WT (black) and *efgA* (blue). Log2FC of each gene in response to formaldehyde (5 min) is plotted.

### Investigating known modulators of formaldehyde tolerance

We wanted to understand the potential roles of genes we previously found to be implicated as relevant to the cellular response to formaldehyde. These genes were identified in one of three ways: (1) genes for which mutations were identified that led to increased formaldehyde resistance and growth on formaldehyde as a sole substrate, (2) the formaldehyde stress response of methanol-grown stationary-phase cultures to initiating growth in methanol supplemented with 4 mM formaldehyde resistance, or (3) the small number of genes with differential expression in a population with high phenotypic tolerance to formaldehyde [28,50].

The first comparison involves beneficial mutations identified when *M. extorquens* was previously evolved in the laboratory to use formaldehyde as a growth substrate. In these experiments, mutations in *efgA* were the predominant class, however, point mutations in *def* (peptide deformylase)*, ttmR* (MarR-like homolog) [51], and *efgB* (predicted adenylyl cyclase) were found to modulate formaldehyde tolerance as well. To assess the potential role of each of these genes in the wild-type response to formaldehyde, we examined their expression as well as that of a second, distant homolog to *def* (*Mext_3949*) and *efgA* (*Mext_4271*) that are present in the genome (Table S4). With the exception of *efgA* and *efgB*, the expression of all of the genes was unchanged by formaldehyde. By the 5 min timepoint *efgA* expression had increased by 3.7× and by the 180 min timepoint, expression was maximal (>17×) (Figure S6A). Expression of *efgB* increased more gradually: unchanged at 5 min, 1.5× at 20 min, 1.8×at 40 min and also maximal at 180 min (3.9×) (Figure S6B). Interestingly, although the Δ*efgA* mutant had an overall subdued response when compared to wild type, changes in *efgB* expression were observed much sooner; 2.5× at 5 min, 3.1× at 20 min and subsiding by 40 min (2.0×). These data demonstrate that *efgA* and *efgB* are upregulated in response to formaldehyde and that the temporal response of *efgB* is sped up in the absence of EfgA.

The second comparison we made was between the formaldehyde stress response of mid-exponential phase succinate-grown cells and methanol-grown stationary phase cells, as was used previously [50]. In the previous experiment, transcriptomics was performed four hours after formaldehyde exposure by inoculation of stationary phase cells into fresh medium; in these conditions, formaldehyde toxicity leads to death of the majority of the cells within the population. In this ‘death phase’, ~54% of genes were significantly up- or down-regulated. (Log2FC>1, padj<0.001). To determine whether the formaldehyde stress response was conserved between treatments, we compared trends in gene expression between them, using only genes that were differentially expressed in mid-exponential succinate-grown (Figure S7). Overall, there was a positive correlation between the two responses. There were expression changes that were anti-correlated, however, given that the fates of cells in these two treatments are so distinct (quiescent versus death), it was unsurprising to find at least some differences.

The third comparison was between the formaldehyde stress response studied here and the limited number of genes identified as DEGs when selection for high phenotypic tolerance within wild-type *M. extorquens* resulted in a population with an elevated spectrum of formaldehyde tolerance [50]. Transcriptomic analysis of a subpopulation with increased formaldehyde tolerance found that tolerant cells had a distinct transcriptomic profile that consisted of 24 differentially expressed genes. To determine if these genes had a general role in the wild-type response to formaldehyde, we examined their expression in both treated and untreated conditions (Figure S8). While about a third of the genes were unchanged in the untreated and the formaldehyde-treated conditions, the rest were either differentially expressed in both conditions or only upon formaldehyde treatment. In instances where genes were differentially expressed in both conditions, the Log2FC were much larger in the formaldehyde-treated samples.

### In the absence of EfgA cells have increased response to proteotoxicity and genotoxicity

Thus far, our analyses showed that in both genotypes there is a strong transcriptional response to formaldehyde stress. To understand the beneficial role of EfgA, we sought to examine which aspects of the transcriptome were changed in the Δ*efgA* mutant upon formaldehyde exposure but not in the wild type. We reasoned that, in the absence of the standard formaldehyde response machinery, the cells might more severely experience the stress that EfgA would typically shield them from. Therefore, we compared the expression fold-changes between the two genotypes at each of the early timepoints (5, 20, 40, 180 min) and extracted the genes that were shown to be differentially expressed only in the Δ*efgA* mutant at any of these times. For those genes, we investigated the full temporal response in both genotypes. From these analyses, three patterns emerged: First, in many instances a similar expression change occurred in both genotypes but occurred much earlier in the Δ*efgA* mutant. Thus, these expression patterns were analogous to that of *efgB*, described above. Second, there were instances where the expression of a gene was only different in the Δ*efgA* mutant. Third, and least common, there were instances where the genotypes elicited opposite expression changes for a given gene.

We hypothesized that genes that were differentially expressed in the *efgA* mutant but not WT would be indicative of formaldehyde-induced stress experienced when EfgA was not present. We performed enrichment analyses on this set of 212 genes (134 upregulated, 78 downregulated) (Figure 11). Although fewer categories emerged compared to WT, again categories related to protein homeostasis and DNA Metabolism & Repair were highly represented in the upregulated genes. Examination of those in the Protein Folding category of the *efgA* mutant and the Heat Shock category of WT showed that even when both strains were upregulated for a particular gene, the *efgA* mutant generally showed more significant increases in expression (Figure 10A). Considering that the *efgA* mutant has a subdued global transcriptional response compared to WT, these data suggest that the *efgA* mutant may experience more potent protein stress despite the greatly muted physiological response in terms of growth. We similarly examined the two DNA-related categories upregulated in the *efgA* mutant (Nuclease activity and DNA metabolism & repair). Between the two categories we identified 12 unique genes and again found that the *efgA* mutant generally showed larger increases in expression among them (Figure S9). Notably, these genes do not overlap with those of the WT DNA recombination/transposition category, which were nearly all more significantly upregulated in WT compared to the *efgA* mutant (Figure 10B). It is possible that in the *efgA* mutant, where cell growth and DNA replication continued despite toxic levels of formaldehyde, there is more extensive DNA damage experienced than is sustained by WT under the same conditions.

**Figure 11.**
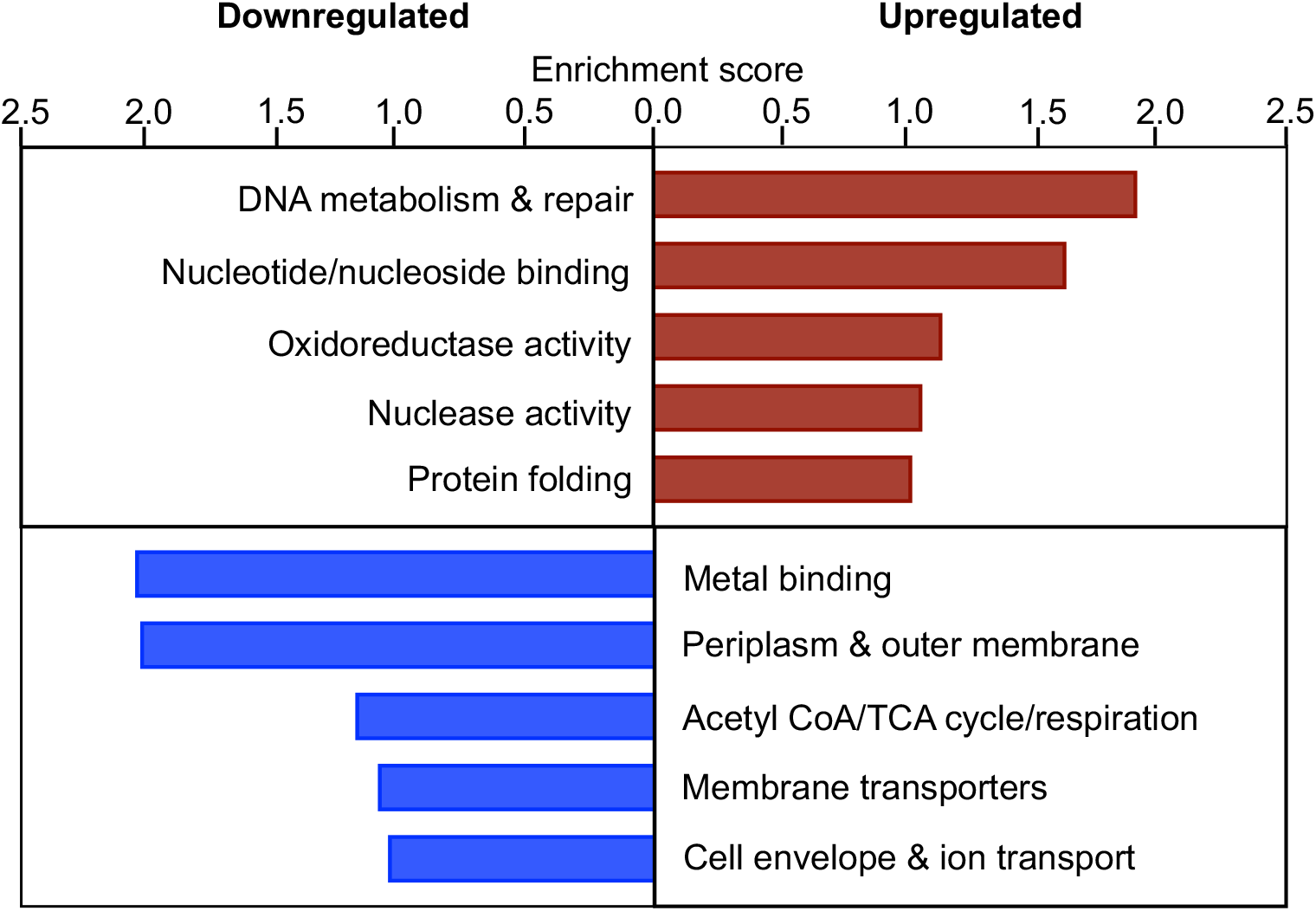
Gene ontology of the early formaldehyde response of Δ*efgA* mutant. Summary of functional group response of Δ*efgA* mutant at 5min (fastest response) and 20min (most intense response) was obtained using DAVID 6.7. Enrichment scores for each functional group are plotted for upregulated (red) and downregulated genes (blue).

## DISCUSSION

Formaldehyde generation is intrinsic to metabolism in all domains of life. In methylotrophs, formaldehyde is generated through high flux methanol utilization pathways and is present intracellularly in substantial amounts during growth on methanol. In *M. extorquens*, the formaldehyde oxidation pathway is the first line of defense for excess formaldehyde as it naturally detoxifies formaldehyde [25]. In such a system, formaldehyde stress, and perhaps even accumulation, are likely to occur upon particular metabolic perturbations. However, the global response to formaldehyde stress is unknown. By characterizing the temporal transcriptional response to exogenously supplied formaldehyde we gained critical insight into the formaldehyde stress response of *M. extorquens* and the role that EfgA, a recently identified formaldehyde sensor, plays in that response.

In wild type, the response to formaldehyde is rapid and immense. For this work, we opted to treat early exponential phase, succinate-grown cells with 5 mM formaldehyde. In these conditions, where formaldehyde is not generated as a central carbon intermediate, we anticipated that cells would not be ‘primed’ for formaldehyde exposure. Still, the response was rapid; by our earliest timepoint (5 min), 1,783 genes (37% of genome) were differentially expressed in response to formaldehyde. Notably included in this early response was a significant downregulation of genes encoding methanol dehydrogenase and the last step in the formaldehyde oxidation pathway, catalyzed by Fhc. As formaldehyde is typically generated by methanol dehydrogenase during growth on methanol, it stands to reason that the response represented negative feedback to cut off excess formaldehyde at the source. Alternatively, a decrease in Fhc activity potentially contributes to eliminating excess formaldehyde through accumulation of methenyl-dH_4_MPT, known to increase Fae activity (i.e., consumption of free formaldehyde) by an unknown mechanism [27]. Beyond one-carbon pathways, wild-type induces a number of genes involved in protein and DNA stress, both of which could be expected as formaldehyde has numerous targets to invoke damage.

Formaldehyde cytotoxicity has typically been attributed to the damage it causes to DNA. In many organisms, such as *Salmonella enterica* and *Escherichia coli*, but this simple interpretation is confounded by conflicting data [52–56]. Inconsistent phenotypes have also been reported in formaldehyde exposure studies used to understand the tolerance of human cell lines to DNA-protein crosslinks [57–61]. Furthermore, formaldehyde is also toxic in non-dividing neural cells that should be tolerant to transient DNA-protein crosslinks [62,63]. In one study, researchers challenged the notion that formaldehyde cytotoxicity was due to DNA damage and demonstrated that formaldehyde was in fact a proteotoxic stressor that caused the accumulation of misfolded proteins across various human cell lines [64]. In the wild-type response, upregulation of genes involved in DNA recombination and transposition included *recA*, a gene typically associated with the SOS response. The SOS response is a bacterial response to DNA damage that simultaneously repairs DNA damage and can produce adaptive mutations as a means of survival [65] while homologous recombination has been shown to be important for resolving formaldehyde-induced DNA-protein crosslinks [58]. This suggested that part of the wild-type response to formaldehyde aims at reversing formaldehyde-induced DNA damage. Notably, in the Δ*efgA* mutant, the SOS response was also muted; however, two distinct groups of genes associated with DNA metabolism were enriched for in that strain. These upregulated genes included nucleases and DNA repair proteins and in wild type, the expression of these genes was nearly unchanged. This finding suggested that the Δ*efgA* mutant failed to activate the SOS response and likely experienced DNA damage that wild type did not. Further, as the Δ*efgA* mutant was actively dividing, even after formaldehyde treatment, we suspect that there was a distinct burden on the Δ*efgA* mutant to maintain the integrity of its genome.

By including both the wild type and the Δ*efgA* mutant, lacking the recently identified formaldehyde sensor EfgA, we found that the observed global response is largely mediated by EfgA. EfgA is predicted to induce translational arrest in the presence of elevated formaldehyde and thus, protect cells from formaldehyde-induced protein damage [28,51]. We hypothesized that cells lacking EfgA would show signs of increased protein stress such as upregulation of chaperone and heat shock proteins. We did not anticipate, however, that cells lacking EfgA would have a dampened response to formaldehyde altogether, only changing expression of ~37% as many genes as wild type. Additionally, the magnitude of the fold changes for differentially expressed genes was overall decreased and the Δ*efgA* strain returns to its pretreatment state even when formaldehyde is still present. Notably, despite the lessened global response to formaldehyde, the Δ*efgA* mutant exhibited stronger signs of increased protein stress, when compared to wild type. In both strains, gene ontology analyses indicated that protein folding/stress genes were upregulated but significantly moreso in the Δ*efgA* mutant. Together, these data support our initial hypothesis of heightened protein stress but also suggest that the role of EfgA extends beyond protecting the proteome. It is further possible that EfgA may mediate a transcriptional response to formaldehyde via its impact on translation, as transcriptional regulation can certainly be linked to translation [66,67].

The difference between EfgA-formaldehyde-mediated translational arrest and that of kanamycin, a known translational inhibitor, was stark. In comparing the response of wild type and the Δ*efgA* mutant to kanamycin, there were no notable differences between genotypes which further confirmed that the impact of EfgA is specifically dependent on formaldehyde. In both strains, significant death occurred with kanamycin treatment. Curiously, death preceded the transcriptional response. By 40 min, even when 10% of the population had already died, there was essentially no population-level transcriptional response. Eventually a distinct transcriptomic profile emerged which bore some overlap with translational inhibition by EfgA-formaldehyde. These differences emphasized the significantly different timescales of the transcriptional response to EfgA-formaldehyde versus kanamycin but showed that there were commonalities, including enriched downregulation of genes related to motility and signal transduction pathways. This response may stem from cells attempting to conserve energy under the heavy stress cessation of translation imposes. One possible explanation for the lack of an early kanamycin response is that cells do not have a mechanism for rapid, uniform detection of kanamycin as they do for formaldehyde. As a minority of cells begin to experience kanamycin stress and succumb to death, their transcriptional response may be overshadowed by the bulk population. When a distinct transcriptional response to kanamycin eventually emerged, it displayed enriched upregulation of translation-related functions and downregulation of other cellular metabolic processes. Intriguingly, despite the rapid cessation of translation provided by EfgA under formaldehyde stress, we do not detect enrichment of translation-related genes in wild-type exposure to this stressor.

Herein, we demonstrate that *M. extorquens* has a rapid, coordinated response to formaldehyde stress that is largely mediated by EfgA. This response induces growth arrest; however, arrested cells maintain metabolic activity and degrade free formaldehyde present in the environment. The arrested cells also launch a coordinated transcriptional response that targets genes to i) minimize the biosynthesis of free formaldehyde, ii) increase formaldehyde consumption by Fae, and iii) mitigate DNA and protein toxicity. In the absence of EfgA, cells continue to grow in spite of exogenous formaldehyde and by comparison to wild type have a lessened but still impressive response to formaldehyde. We hypothesized that if EfgA protects the cells from formaldehyde-induced stress, it is likely that bypassing the EfgA-mediated response would negatively impact cells. The decreased growth rate of the Δ*efgA* mutant in the presence of formaldehyde supported this hypothesis. In the absence of EfgA the transcriptional response that is unique, including the upregulation of genes involved in DNA/protein stress and the downregulation of genes involved in cell envelope metabolism, transport, and metal binding, suggest that the mutant experiences increased formaldehyde stress. Thus, we conclude that in addition to EfgA arresting growth and translation, it also functions directly or indirectly as a sizable contributor to the transcriptional formaldehyde stress response in *M. extorquens*.

---

**Figure S1.**
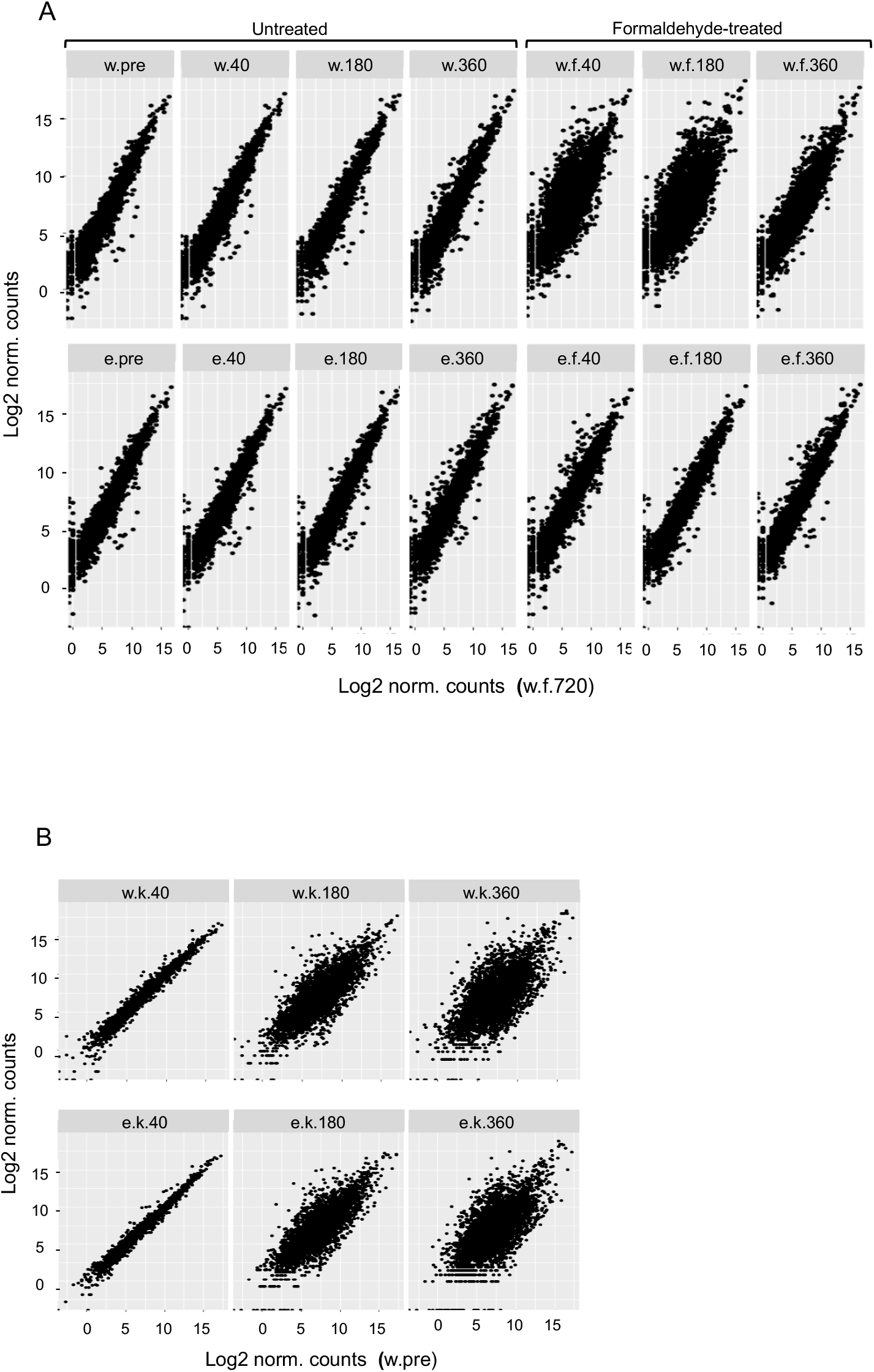
Single replicate screen of transcriptional response identifies samples of interest. A) Single replicate data for RNA-Seq of WT (‘w’, top panel) and Δ*efgA* mutant (‘e’, bottom panel) of *M. extorquens* grown in succinate minimal medium and untreated or treated with 5mM formaldehyde (‘f’) for 0 (‘pre’), 40, 180, 360, 720 min. Log2 transformed counts of the entire genome for each sample were plotted against those of WT formaldehyde-treated 720 min post treatment (w.f.720, x-axis) to establish the timescale of the transcriptomic response and identify the conditions most similar to the 720 min timepoint. B) Single replicate data for RNASeq of WT (‘w’, top panel) and Δ*efgA* mutant (‘e’, bottom panel) of *M. extorquens* grown in succinate minimal medium and treated with 50 μg/mL kanamycin (‘k’) for 40, 180, 360 min. Log2 transformed counts of the entire genome for each sample were plotted against those of WT pretreatment (w.pre, x-axis). Each point represents one gene.

**Table S3. Genes differentially expressed in untreated samples.**

Genes exhibited temporal expression changes in the absence of a stressor. To identify genes differentially regulated during the timecourse in untreated samples, the log2 of normalized counts (of each genotype at each timepoint) was compared to the WT pretreatment sample. Values highlighted in gray lack statistical significance (padj >0.001).

**Figure S2.**
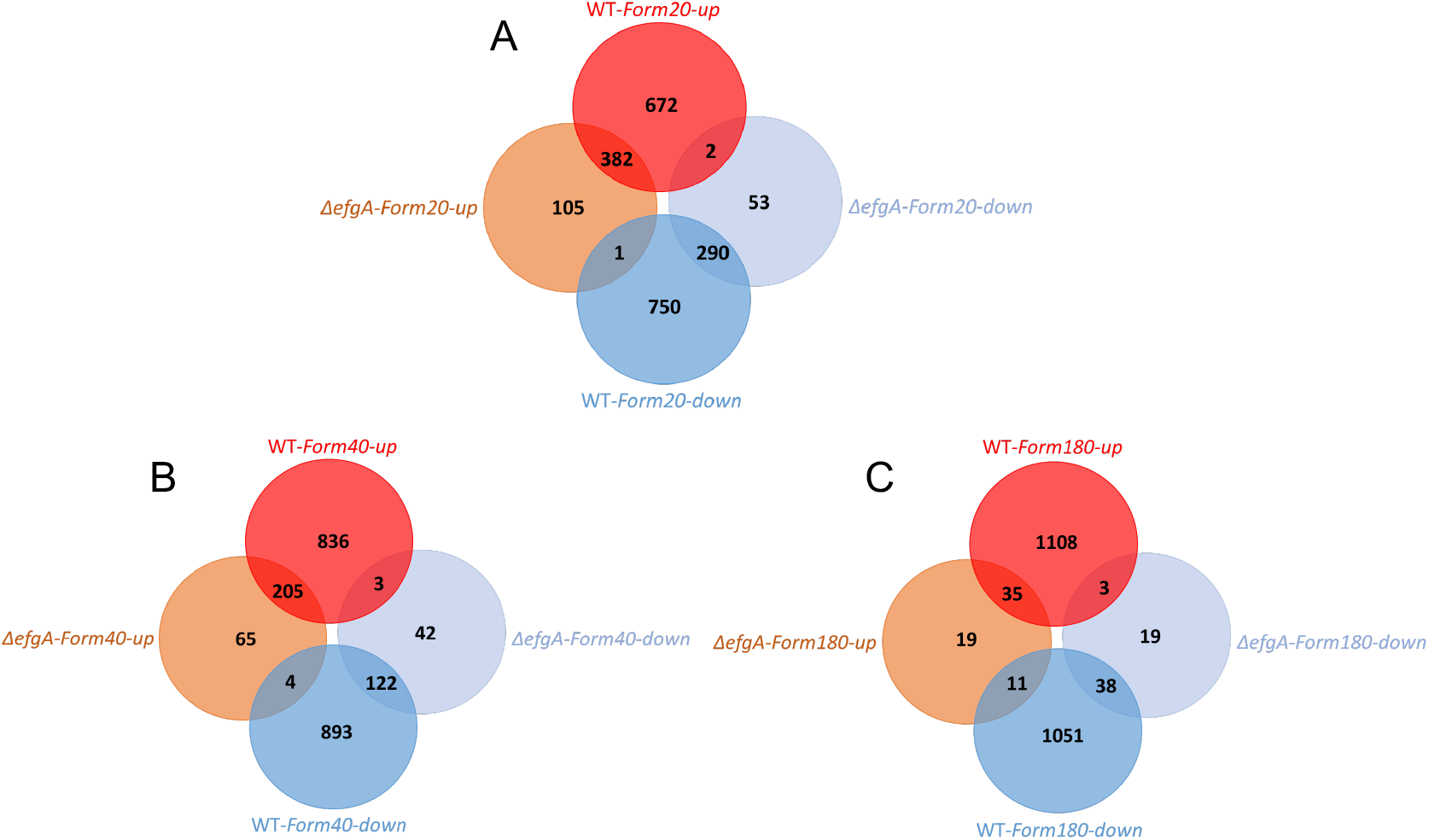
Venn diagram of genes differentially expressed in WT and Δ*efgA* mutant upon formaldehyde treatment. Each diagram represents a different timepoint A) 20 min, B) 40 min, C) 180 min. Genes differentially expressed (|log2-fold change| >1) were identified by dividing Log2FC of sample/Log2FC of pretreatment sample (of the respective genotype). Venn diagram for the earliest (5 min) timepoint is depicted in Figure 5.

**Figure S3.**
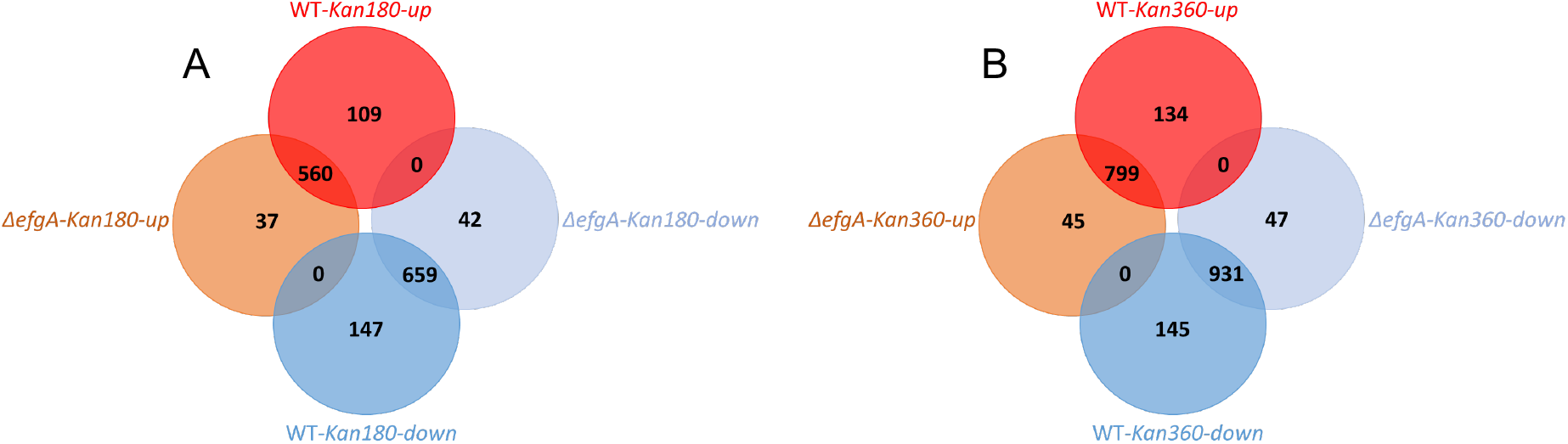
Response to kanamycin is similar in WT and Δ*efgA* mutant. Venn diagrams of genes differentially expressed in WT and Δ*efgA* mutant upon kanamycin treatment. Each diagram represents both genotypes at a distinct timepoint: A) 180 min and B) 360 min. Genes differentially expressed (|log2-fold change| >1) were identified by dividing Log2FC of sample/Log2FC of pretreatment sample (of the respective genotype).

**Figure S4.**
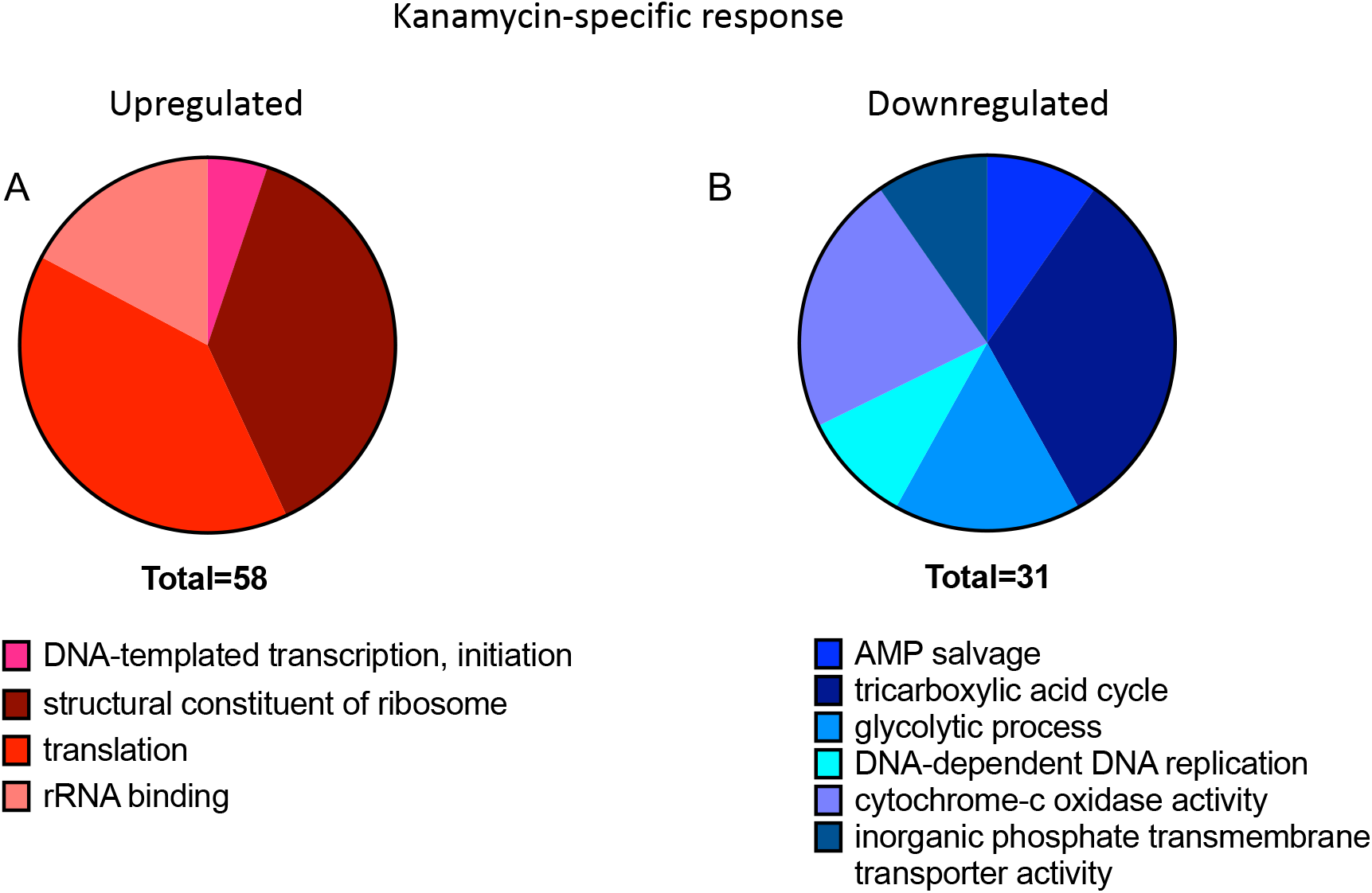
Gene enrichment of the kanamycin-specific response. Summary of functional group response of WT to kanamycin (360 min) was obtained using Comparative GO. All functional groups shown had >2 genes and p<0.05.

**Figure S5.**
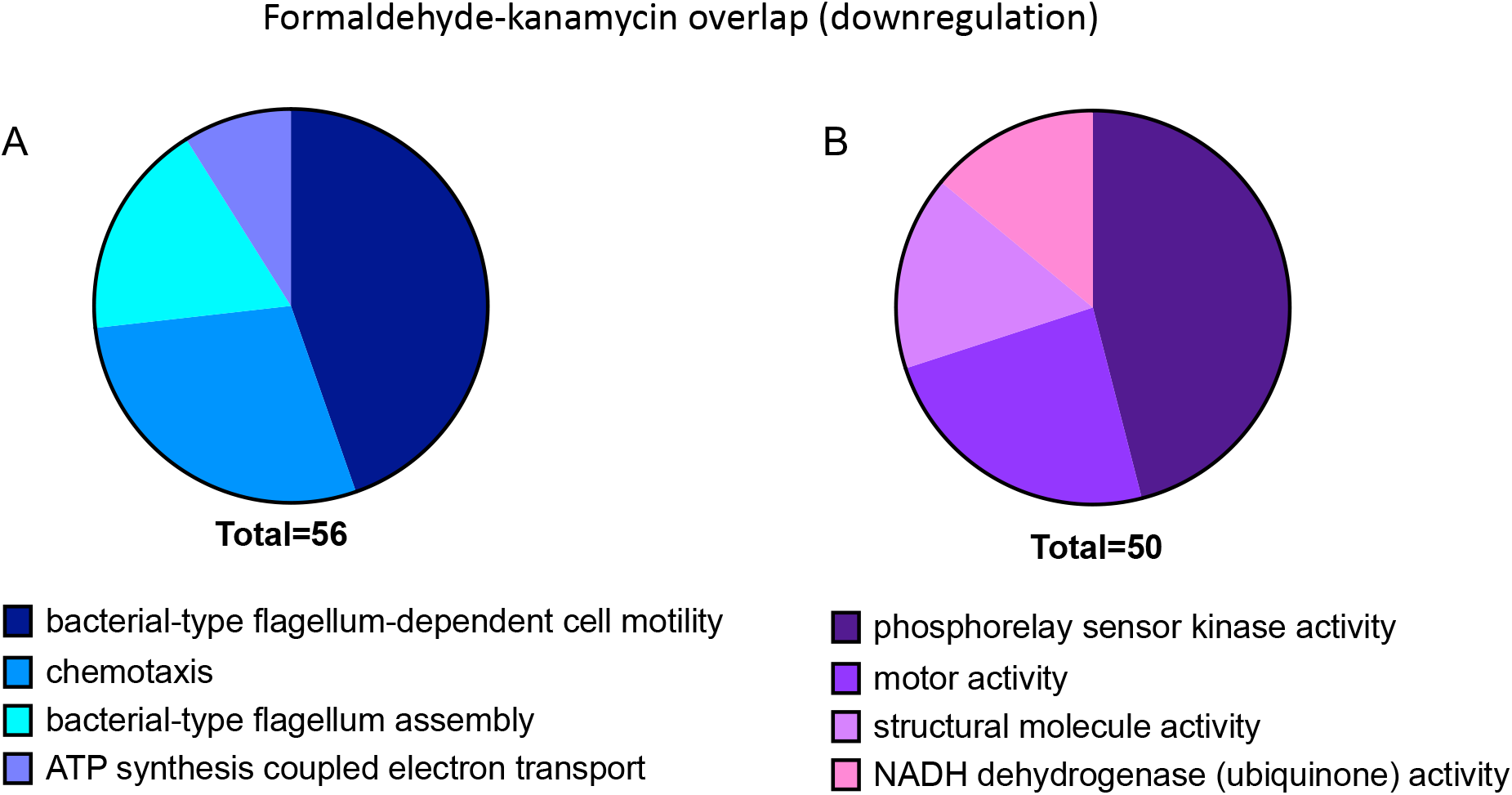
Enrichment of downregulated genes in both the kanamycin and formaldehyde responses. The overlap of gene functional groups in the downregulated responses of WT to kanamycin (360 min) and formaldehyde (5 min) was obtained using Comparative GO. Gene function is categorized by A) Biological Process B) Molecular Function. All functional groups shown had >2 genes and p<0.05.

**Table S4. Changes in expression of genes identified during evolution to growth on formaldehyde.**

For each genotype, the first column shows the Log2FC (at 5 minutes) compared to its pre-treatment condition, the second column shows the Wald test statistics and the third column shows the FDR-adjusted p-values. Close homologs to *efgA* and *def* were included due to their uncertain biological role. Only *efgA* in WT and *efgB* and the *efgA* homolog in Δ*efgA* showed significant changes.

**Figure S6.**
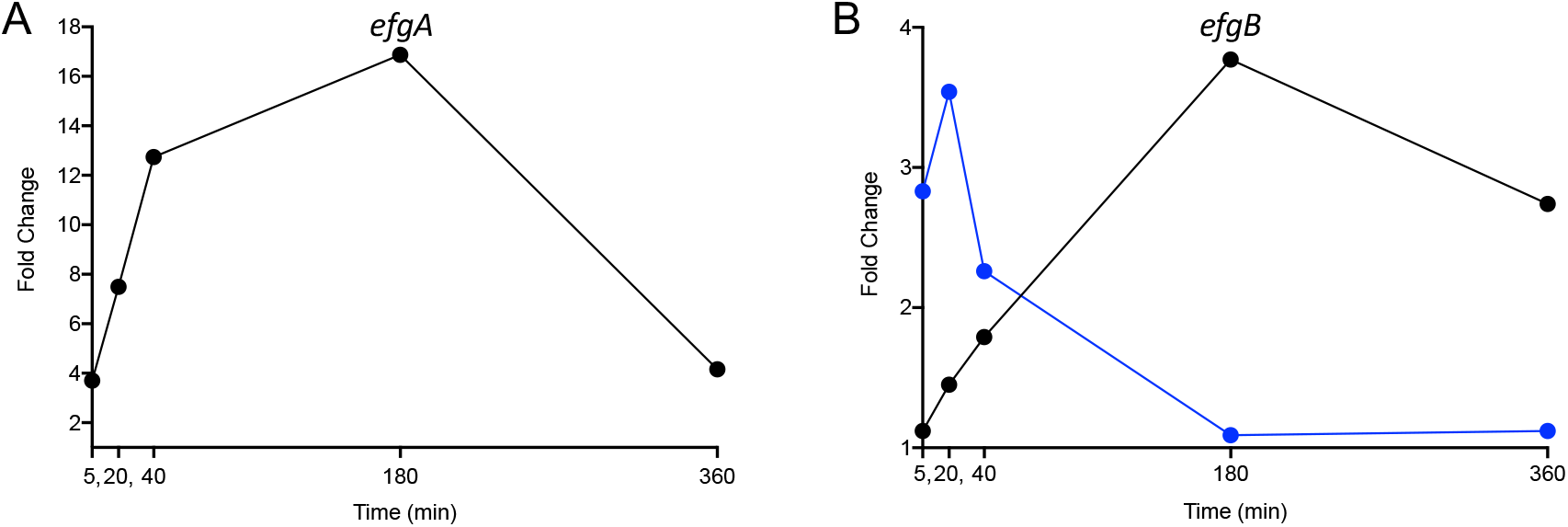
*efgA* and *efgB* are upregulated in response to formaldehyde. Temporal expression of A) *efgA* in WT and B) *efgB* in WT (black) and the *efgA* mutant (blue) in response to formaldehyde treatment. Log2FC of each gene is plotted.

**Figure S7.**
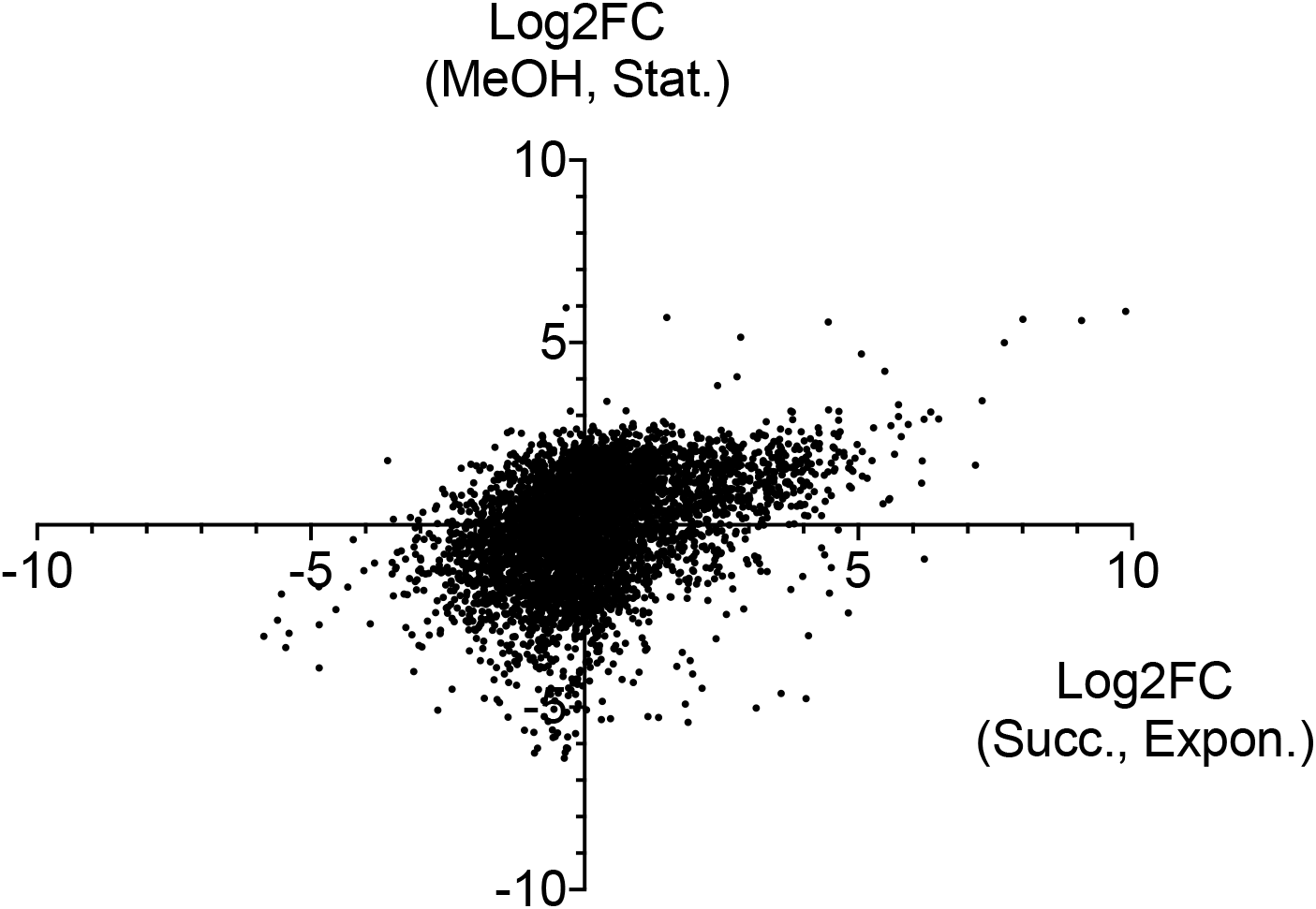
Comparison of early formaldehyde response in different growth conditions. Expression changes (Log2FC) of all genes at the earliest timepoint (5 min) here, where succinate-grown cells were treated with 5 mM formaldehyde in early exponential phase of growth are plotted against expression changes of the same genes from a previous experiment [50] where methanol-grown cells were treated with 4 mM formaldehyde in stationary phase of growth and assessed at 4 hr.

**Figure S8.**
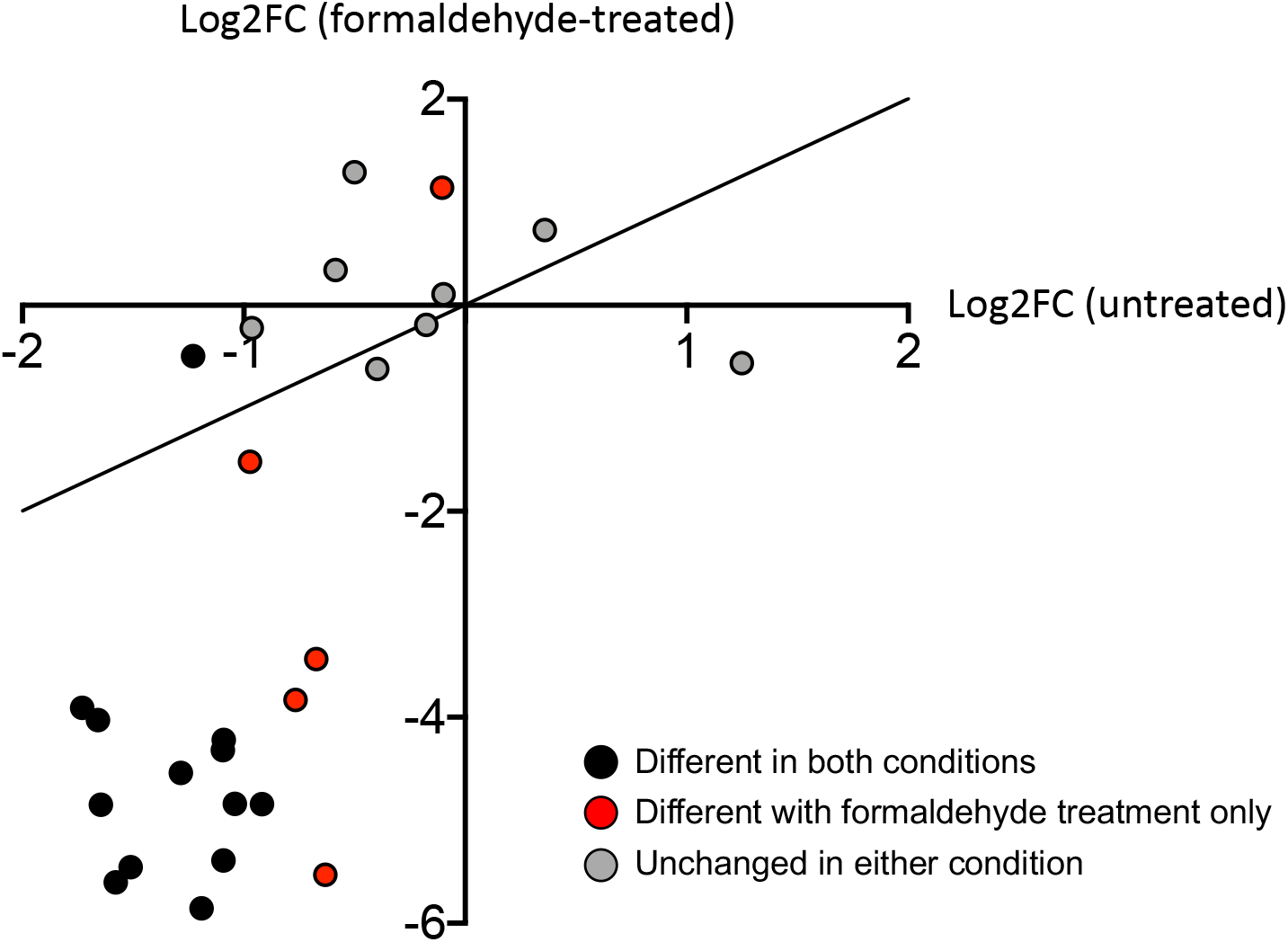
Response of genes involved in formaldehyde tolerance heterogeneity. Expression changes (Log2FC) of genes predicted to be differentially expressed in formaldehyde tolerant subpopulations were examined in untreated (x-axis) and formaldehyde-treated (y-axis) samples.

**Figure S9.**
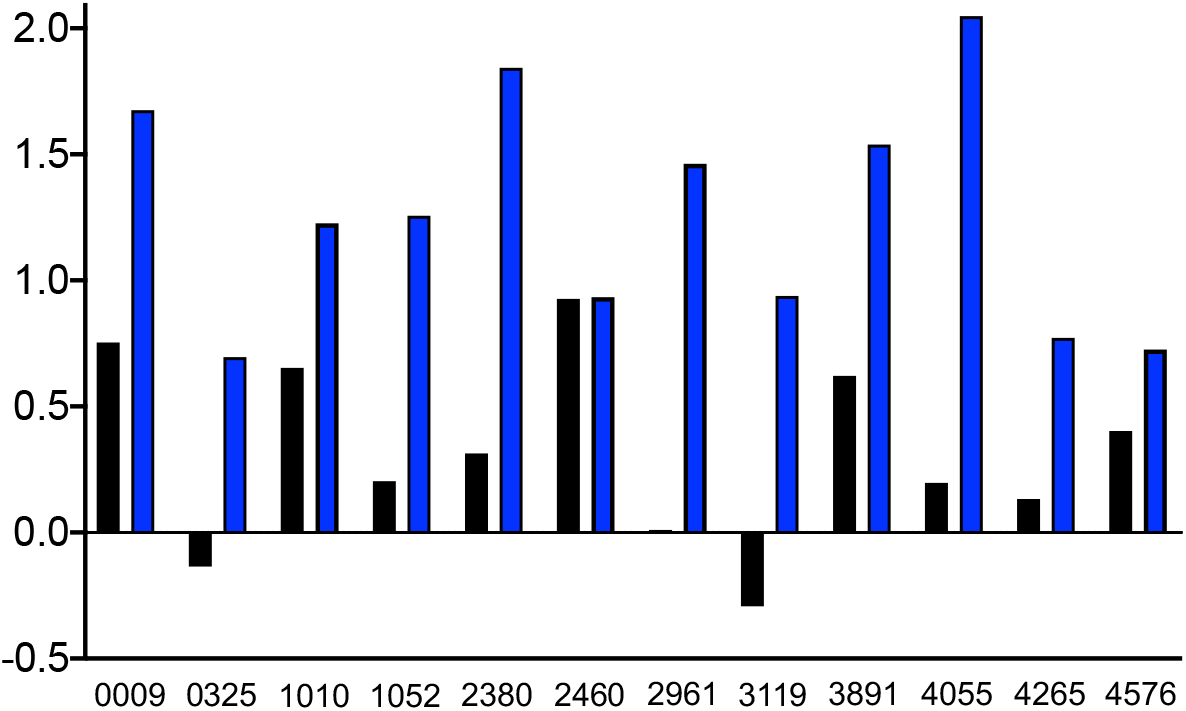
Genes involved in Nuclease activity and DNA metabolism & repair suggest the Δ*efgA* mutant experiences different DNA stress than WT. Expression of genes involved in nuclease activity and DNA metabolism & repair are shown for WT (black) and *efgA* (blue). Log2FC of each gene in response to formaldehyde (5 min) is plotted.

## Acknowledgements

We thank members of the Marx and Bazurto laboratories for critical reading of this manuscript and Jessica A. Lee for assistance with conducting experiments. Christopher J. Marx and Jeffrey E. Barrick were supported by funding from the Army Research Office grant W911NF-12-1-0390. Siavash Riazi was supported by fellowships from the Bioinformatics and Computational Biology Graduate Program and the Department of Biological Sciences. Support was also received via pilot grants to Jannell V. Bazurto from the BEACON Center for Evolution in Action (parent award DBI-0939454) and Christopher J. Marx from the Institute for Modelling Collaboration and Innovation (parent award P20GM104420).

